# Sexual dimorphism and biomechanical loading in occipital bone morphological variation

**DOI:** 10.1101/2022.04.05.487198

**Authors:** Kara C. Hoover, Geoffrey P. Thomas

**Affiliations:** Department of Anthropology, University of Alaska, Fairbanks Alaska 99775; Department of Anthropology, Florida State University, Tallahassee Florida 32306

**Keywords:** occipital bone morphology, sexual dimorphism, biomechanical loading, subsistence, activity patterns

## Abstract

We explore the contribution of biological sex and biomechanical activity from subsistence to occipital bone variation. Previous studies have used occipital bone traits to determine biological sex and identify ancestry to differing degrees of success. Biomechanical stress from variation in subsistence and gender-based divisions of labor could perhaps explain some of the noise in the signal for these grouping variables. To explore this possibility, we used metric (foramen magnum length and breadth, external occipital protuberance depth, lambda-inion length, bicondylar breadth) and nonmetric traits (general occipital form, presence of a nuchal crest, and nuchal line count). We collected original data and mined published data for our analysis using skeletal collections of Native American hunter-gatherers and horticulturalists, a historic military site, and contemporary study collections. We find that the foramen magnum area exhibits sexual dimorphism and is not influenced by subsistence but the accuracy of sex estimation is only 71%, suggesting the chance of being correct at slightly more than two-thirds. All traits exhibited sex-based variation but only bicondylar breadth and lambda-inion exhibited subsistence-based variation. Given the limited amount of variance explained by either sex or sex and subsistence, biomechanics may still play a role but not from the influence of subsistence practices. Additional data from a wider array of skeletal samples, perhaps with known occupation, is warranted if we are to understand how occipital variation is shaped.

**PRACTITIONER POINTS:**

- The study examines the role of biological sex and biomechanical stress in shaping occipital morphology.
- We introduce an area variable to describe the foramen magnum and apply ectocranial metric and nonmetric states for the first time to sex estimation.
- We share methods and R code for cleaning data, testing assumptions, and analysis of research questions.

## INTRODUCTION

Occipital bone morphological traits have been used to estimate biological sex (Gapert, Black, & Last, 2009; Gapert & Last, 2008; Giles & Elliot, 1963; Gulekon & Turgut, 2003; Holland, 1986; Hsiao, Chang, & Liu, 1996; Macaluso, 2011; Sholapurkar, Virupaxi, & Desai, 2017; Uysal, Gokharman, Kacar, Tuncbilek, & Kosar, 2005; Williams, 1987; Zdilla, Russell, Bliss, Mangus, & Koons, 2017) and ancestry or temporal change within a group over decades (Moore-Jansen, 1989; Wescott, 1996; Wescott & Moore-Jansen, 2001; Williams, 1987). Ancestry estimation or population affinity (often considered euphemisms for race) is critiqued in biological anthropology as a method that, at its core, contradicts a well-established fact that race has no biological basis—no trait is discrete to a population, however defined (DiGangi & Bethard, 2021). Sex estimation has not shared the same level of sustained controversy but current work is questioning the limitations of reducing the concepts of sex and gender to estimates from skeletal elements based that, among other things, may overlook constructions of gender that are not linked to sex (Moral, 2016; Zuckerman, 2020). The lack of consistency in results for sex and ancestry estimation in previous studies of the occipital may be explained by another source of variation, biomechanical activity that might be impacted by gender-based division of labor and subsistence. We examined occipital bones from human remains belonging to Native Americans, a historic military cemetery, and contemporary study collections to explore this possibility. We had two datasets. The Original Dataset generated for this study used Florida-based collections (Windover Mortuary Pond, Hutchinson Burial Mound, 19^th^ Century St. Mark’s military cemetery, and a modern study collection). The Original Dataset variables include three nonmetric traits and five metric traits (three describing the base of the occipital bone and two describing the morphology of the bone surface). The Mined Dataset consists of published data for horticultural Native American skeletal assemblages found in South Dakota and modern study collections. The Mined Dataset is not analyzed but is combined with the Original Dataset, resulting in a Combined Dataset consisting of three metric variables for the base of the skull that are common across all studies.

Changes to the cranium (particularly the forward placement of the foramen magnum) are significant in human evolutionary history due to human obligate bipedalism (Ross & Ravosa, 1993; Russo & Kirk, 2013, 2017). Sexual dimorphism of the skull has a similarly long evolutionary history and has been noted in our early biped ancestors (Kimbel & Rak, 2010). The occipital bone features a series of traits linked to sexual dimorphism in two main regions, the basicranium and the thick area surrounding the external protuberance. The posterior portion of the occipital exhibits muscle attachment sites along the nuchal region for head movement. Because the bone is thick and therefore more resistant to taphonomic processes, it has been the subject of forensic investigation as a potential diagnostic tool in sex assessment of unidentified and fragmentary or damaged human remains (Gapert & Last, 2008; Gulekon & Turgut, 2003; Hsiao et al., 1996; Sholapurkar et al., 2017; Wescott & Moore-Jansen, 2001).

The basicranium is considered the more naturally sexual dimorphic area of the skull with two traits, the foramen magnum and the occipital condyles. The foramen magnum of females tend to have smaller lengths and widths than males (Catalina-Herrera, 1987; Holland, 1986; Zdilla et al., 2017) and the condylar regions of females have shorter bicondylar breadths than males (Gangrade, Saini, Yadav, & Vyas, 2013; Holland, 1986; Kunar & Nagar, 2014; Macaluso, 2011; Sholapurkar et al., 2017). These size differences may be due to constraints exerted by overall sex-based differences in cranial size (Zdilla et al., 2017; S. Zuckerman, 1955)—male heads tend to be larger and heavier than females, but not always. The basicranium is fully developed by age 7 and the foramen magnum does not remodel later in life for two reasons: the nervous system is developed in early childhood and it has no biomechanical action in its functional role in providing passage from the brain to the body for the medulla oblongata, cranial nerves, and blood supply (Scheuer & Black, 2000). Thus, the foramen magnum, and any sexual dimorphism exhibited by it, reflects early life developmental trajectory. The other region of the basicranium (the occipital condyles), however, may be altered in life through biomechanical processes during articulation with the superior facets of the first cervical vertebra for head rotation, flexion, and extension. Studies show that condyle size asymmetry may result from handedness (Uysal et al., 2005) which further suggests biomechanical pressures acting on the area. As a result, sex-based differences in this region (Gangrade et al., 2013; Holland, 1986; Kunar & Nagar, 2014; Macaluso, 2011; Sholapurkar et al., 2017) may well be a function of activity rather than inherent biological difference between the sexes (Zdilla et al., 2017; S. Zuckerman, 1955) and that may explain why occipital condylar metrics have not been consistently found to have a strong degree of separation between the sexes (Gapert et al., 2009; Wescott & Moore-Jansen, 2001).

The posterior ectocranial portion of the occipital bone also exhibits sex-based morphological variation. It is characterized by heavy buttressing and serves as an attachment site for major neck muscles. The superior nuchal line is generally well developed in adults and has a crest at the center, called the external occipital protuberance (White, 1991). The splenius capitis for head extension and the trapezius for scapula movement and arm support attach to the superior nuchal line (Standring, 2015). Attaching both to the superior and highest nuchal lines are the occipitalis and the epicranial aponeurosis muscles that control scalp movement (Standring, 2015). The crest that marks the external occipital protuberance is the site of the nuchal ligament attachment, which supports the weight of the head (Standring, 2015) and may be particularly useful in interpreting activity patterns due to the association between well-developed nuchal ligaments and running (Swindler & Wood, 1973). Attaching to the inferior nuchal line are a series of muscles used during flexion and extension. The obliquus capitis superior enables head extension and flexion (Standring, 2015). The rectus capitis posterior major is used for both medial and some lateral extension and rotation (e.g., head nodding and wagging) (Standring, 2015). The rectus capitis posterior minor is also used for head extension while serving to restrict movement toward the spinal cord (Standring, 2015). In human males, the nuchal area of the skull is more pronounced than in females (Ebraheim, Lu, Biyani, Brown, & Yeasting, 1996; Gulekon & Turgut, 2003; Hsiao et al., 1996; Olivier, 1975) but no one has examined sex-based differences in activity relative to this region. One final trait that exhibits sexual dimorphism is the length from the lambda (meeting of sagittal and lambdoid suture) to the inion (highest point of the external occipital protuberance), which is longer in females (Olivier, 1975). This trait, however, may be a function of external occipital protuberance morphology because more robust protuberances alter occipital morphology.

As noted above, the history of occipital bone research is strongly biased in favor of evolutionary and forensic inquiries rather than behavioral differences that might either disrupt or increase sexual dimorphism. Occipital traits experiencing biomechanical load may exhibit activity-based variation, which may increase sexual dimorphism or group differences due to workload variation and explain some of the variation previously found in other studies (Marlowe, 2007). To parse traits fixed early in development (under stronger genetic control) from those potentially influenced by biomechanical activity, we contrast the foramen magnum length and breadth to the remainder of traits influenced by muscle use and development (occipital condylar breadth, lambda-inion distance, external occipital protuberance depth, nuchal crest presence, nuchal line count, and general form of the external occipital protuberance). We expect all traits to exhibit sexual dimorphism due to the previously noted association between occipital traits and cranial size and weights, but we expect that ectocranial traits (and possibly bicondylar breadth) may also exhibit variation due to subsistence activities.

## MATERIALS

### Native American Windover Mortuary Pond (8BR246), Hunter-Gatherer Subsistence, Original Dataset (Table 1)

**Table 1:** Materials.

| **Original Dataset (n=98)** | **Subsistence** | **Date** | **n** | **n-female** | **n-male** |
| --- | --- | --- | --- | --- | --- |
| Windover (8BR246) | Hunter-Gatherer | 8,522-7,421 BP<br>Middle Archaic | 47 | 24 | 23 |
| Hutchinson Island Burial Mound (8MT37) | Hunter-Gatherer | 2,000-1,000 BP<br>Late Archaic | 16 | 8 | 8 |
| Modern FSU Teaching Collection | Agriculture | 20th century | 22 | 11 | 11 |
| St. Mark's Military Cemetery (8WA108) | Farming | 19th century | 13 | 0 | 13 |
| **Mined Dataset (n=565)** |  |  |  |  |  |
| Larson (39WW2) | Horticultural | 1679-1733 | 115 | 57 | 58 |
| Leavenworth (39CO9) | Horticultural | 1802-1832 | 30 | 14 | 16 |
| Mobridge (39WW1) | Horticultural | 1600-1700 | 56 | 31 | 25 |
| Hamann-Todd Collection | Agriculture | 1912-1938 | 107 | 54 | 53 |
| Terry Collection | Agriculture | 1899-1941 | 257 | 129 | 128 |
| **Combine Dataset (n=663)** |  |  |  |  |  |

The Florida archaeological site of Windover (8BR246), dates to the Early Archaic period (8,522 −7,421 BP calibrated). The Indian and St. John’s Rivers became population growth hotbeds where the rich environment supported large sedentary base camps with seasonal patterns of small group foraging expeditions to farther-flung resources. The first ‘cemeteries’ emerge at this time in the form of mortuary ponds (Milanich & Fairbanks, 1980) and provide a wealth of information on cultural changes exacted from climate change. The bog was used seasonally for burials and strategically located between the Indian River coastal lagoon system and the St. John’s River—an area rich in marine, fresh water, and terrestrial resources. Growth ring data from mortuary stakes suggest they were harvested in late summer/early fall which points to the period of occupation near the bog (Doran & Dickel, 1988). Residential mobility was limited to a constrained geographic area with limited inter-season travel distances (Adovasio, Soffer, & Page, 2009) away from the bog. The Windover people probably did not fission into smaller groups and most evidence points to the emergence of sedentism (Wentz, 2006). Skeletal materials from this site are unusually well-preserved due to the anaerobic environment of the bog in which they were deposited. Population demography is well balanced between the sexes, and individual ages range from infancy to 65+ (Purdy, 1991). Through the analysis of stomach contents and bone isotopes (Tuross, Fogel, Newsom, & Doran, 1994), the Windover population was found to have a hunter-gatherer subsistence strategy focused on inland riverine, pond, and marsh resources. Of the burials now curated at Florida State University, data were collected on 47 adults (Table 1). The collection is curated by the Department of Anthropology at Florida State University in Tallahassee, Florida.

### Native American Hutchinson Island Burial Mound (8MT37), Hunter-Gatherer Subsistence, Original Dataset (Table 1)

The exact site of the Florida Hutchinson Burial Mound is not known due to initial reporting of the site by a member of the public 1951 and a gap prior to formal archaeological investigation of the area. Well-preserved human skeletal remains were removed from the site by amateur archaeologists, but no funerary objects or other artifacts were recovered. The skeletal remains were studied at Florida State University by Dan Morse. The site is believed to be part of the Hutchinson Island Burial Mound, which was excavated in the mid-1970s and was located on a small cove along the Indian River in Martin County, Florida. The site is described in the original site report as spanning several archaeological periods, prior to European contact through to the 17^th^ century. The large midden underlying the mound is comprised of what appear to be two smaller mounds but is the result of one extending to 250 feet south of the main site, with potential for it to have continued to a large dune along the nearby Atlantic Ocean. Systematic excavations have not taken place and the extent of the site is not fully known. As with Windover, the site is inland but not far from the coast with terrain ranging from coastal dunes to tropical hammock. The analysis of skeletal materials revealed that the teeth were severely worn with few caries (typical of hunter-gatherers and a coarser diet) and several skulls had fractures incurred during life. Three skulls had Inca bones. There were 35 burials found at the main site with 9 mostly complete and the remainder partial (sometimes skulls only and sometimes other bones)—evidence of natural disturbance. There was no patterning to the burials or consistent orientation and no sign of grave preparation (Florida Bureau of Archaeological Research, 1972). Sixteen individuals dating to the Middle Archaic period of Florida were examined for this study. The collection is curated by the State of Florida Bureau of Archaeological Research in Tallahassee, Florida.

### Native American Arikara from South Dakota (multiple sites), Horticultural Subsistence, Mined Dataset (Table 1)

We used published data from a Master’s Thesis (Williams, 1987) on the Northern Plains Arikara populations from three sites along the Missouri River in South Dakota: Leavenworth (39CO9) from 1802-1832, Mobridge (39WW1) 1600-1700, Larson (39WW2) from 1679-1733 (Williams, 1987). These sites date to early in the history of Northern and Great Plains tribal establishments. Mobridge is the earliest of the three sites and was settled prior to contact but all sites overlap with historic contact and the introduction of the horse to the area (Grant, 1995). Populations engaged in mixed economies that included hunting, fishing, crop cultivation, and trade (particularly with the nomadic Sioux) but are perhaps best described as horticultural (Krause, 2016). Cultural homogeneity across Arikara villages was maintained by extensive trade networks that provided opportunities for marriage, adoption, and resulted in direct and indirect contact with Europeans (Grant, 1995). There was a strict gender-based division of labor within households with men engaging in trade and acts of extreme physical exertion (e.g., hunting and warfare/conflict). Women were primarily engaged in farming but also had a wide range of activities (e.g., trade of farmed goods, house building, textile and hide production/processing, and food and tool production) that likely required longer bouts of labor than men (Krause, 2016). There were 201 individuals used in this study: Leavenworth (n=115), Mobridge (n=56), and Larson (n=30) (Table 1). The collection is curated by the Department of Anthropology at the University of Tennessee Knoxville in Knoxville, Tennessee.

### Historic St. Mark’s Military Cemetery (8WA108), Agricultural Subsistence, Original Dataset (Table 1)

St. Mark’s Cemetery (8WA108) was discovered and excavated during road construction in 1965 in Florida (Dailey, Morrell, & Cockrell, 1972). Based on military button patterns, the cemetery was used most likely between 1812-1821. The military post at St. Mark’s was abandoned by 1825 when Spain ceded Florida to the United States. The contents of the site included a total of 16 burials containing human remains (3 are incomplete without crania and one is an incomplete cranium), along with the remnants of toe-pincher cypress coffins fastened together with wrought iron nails with lead heads. The soldiers were likely to have died from malaria while in Fort St. Mark’s infirmary because most of the bodies have no evidence of trauma (other than one broken nose) and were buried in partial uniforms as evidenced by the rarity of uniform buttons and buckles—consistent with being in the infirmary. Soldiers were generally of medium height and suffered many dental problems. Data were collected on 13 adult males (Table 1). The collection is curated by the Department of Anthropology at Florida State University in Tallahassee, Florida.

### Contemporary Study Collection (FSU Teaching Collection), Agricultural Subsistence, Original Dataset (Table 1)

Human remains have been accumulated over a 50-year period with the most common origin from medical schools, although the documentation as to their origins are unknown. The collection consists of mostly older individuals that have been separated by element and therefore represent many co-mingled individuals with isolated crania. Data were collected on 22 adults (Table 1). The collection is curated by the Department of Anthropology at Florida State University in Tallahassee, Florida.

### Contemporary Study Collection (Terry), Agricultural Subsistence, Mined Dataset (Table 1)

We used published data from a Master’s Thesis, shared by Daniel J. Wescott (Wescott, 1996). The Robert J. Terry collection is housed at the Smithsonian Institute, National Museum of Natural History and is comprised of ~1700 unclaimed human remains from local hospitals and morgues from the St. Louis area. Amassed between the years of 1899 and 1941, the collection was compiled after dissection in gross anatomy classes at Washington University. The Terry collection is commonly used as a study collection for anthropological research due to known demographic parameters per individual (e.g., sex, age, race, cause of death, weight) (Hunt & Albanese, 2005). The sample size is 257 (Table 1). The collection is curated by the Smithsonian Museum in Washington D.C.

### Contemporary Study Collection (Hamann-Todd), Agricultural Subsistence, Mined Dataset (Table 1)

We used published data from a Master’s Thesis, shared by Daniel J. Wescott (Wescott, 1996). The Hamann-Todd collection is housed at the Cleveland Museum of Natural History and is comprised of ~3000 unclaimed human remains from morgues and hospitals in the Cleveland area. Amassed between the years 1912 and 1938 by Carl Hamann and T. Wingate Todd, the collection was used in anatomy classes at Case Western University (then Western Reserve University). Hamann-Todd is one of the largest and commonly used study collections for anthropological research due to individual level demographic data (e.g., sex, age, race, cause of death, weight). The sample size is 107 (Table 1). The collection is curated by the Cleveland Museum of Natural History in Cleveland, Ohio.

## METHODS

### Study Inclusion Criteria

Occipital traits have not been found to exhibit age-related morphological differences in adulthood (Holland, 1986; Wescott & Moore-Jansen, 2001). As such we retained all adults with intact bones and did not consider age as a confounding factor in our examination of skeletal sex and subsistence differences.

### Age and Sex Estimation

GT used traditional pelvic and cranial non-metric traits for skeletal sex estimation and a combination of endo/ectocranial suture closure, long bone epiphyseal fusion, and pubic symphysis degeneration methods for age estimation (Brooks & Suchey, 1990; Buikstra, Frankenberg, & Konigsberg, 1990; White, 1991). Age estimates are not available for the mined data.

### Subsistence

The Florida Archaic populations were both hunter-gatherers while the three Arikara sites are best classified as horticultural (see individual site descriptions above). While the Florida St. Mark’s Military Cemetery is from the 19^th^ century and the study collections are from the 20^th^ century, the agricultural subsistence practices in both periods is marked by a diversity of occupations with greater heterogeneity across individuals in terms of their occupations and possible biomechanical stresses.

### Original Dataset Data Collection

Three trials of data collection (GT) were completed using an electronic sliding caliber for five metric traits: depth of external occipital protuberance (Figure 1); distance between the lambda and inion (Figure 2); foramen magnum bicondylar breadth in the sagittal plane, foramen magnum breadth in sagittal plane, and foramen magnum depth/length in transverse plane (Figure 3). The external occipital protuberance is located on the ectocranial midline at the intersection between the occipital and nuchal planes (White, 1991) and was measured for depth at the inion, or center of the external occipital protuberance (Olivier, 1969). The protuberance can be located using the inferior or superior nuchal line as landmarks (Olivier, 1969). Lambda-inion distance is measured from the inion to the point where the lambdoidal suture meets the sagittal suture, the lambda (Olivier, 1969; White, 1991). Maximum bicondylar breadth was measured for distance across the lateral margins of the occipital condyles (Holland, 1986). The internal length and width of the foramen magnum was measured along the sagittal and transverse planes (Buikstra & Ubelaker, 1994; Holland, 1986).

**Figure 1:**
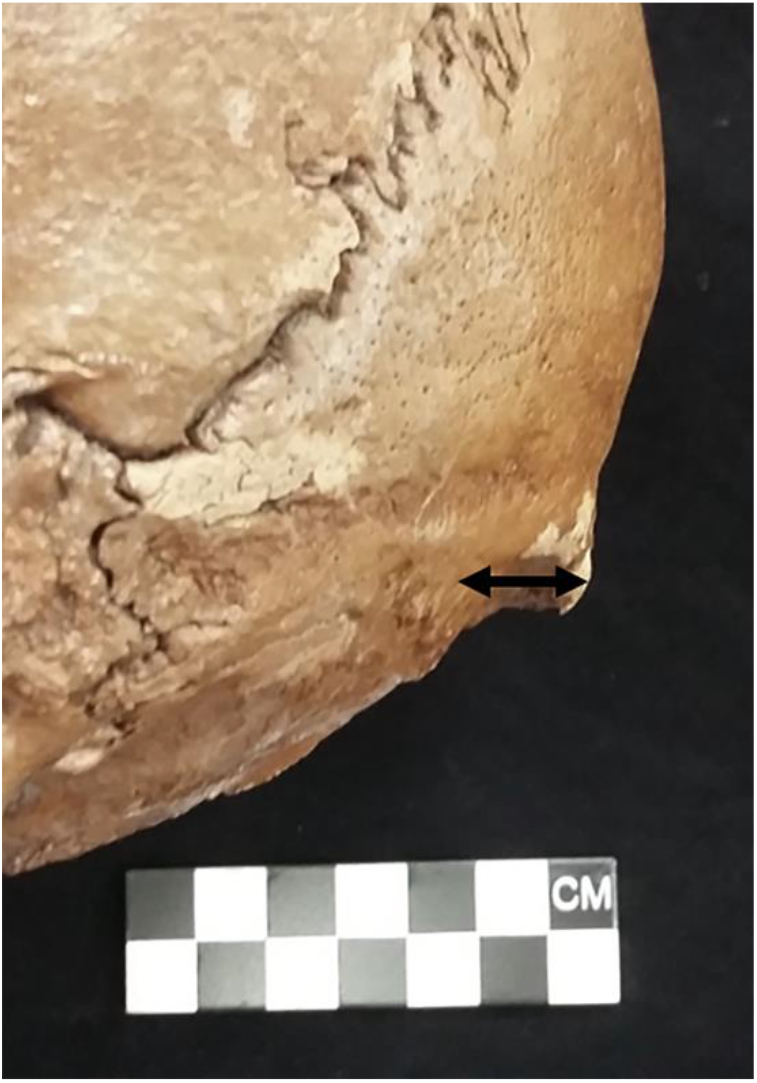
Depth of the external occipital protuberance (Image taken by GT Historic St. Mark’s Military Cemetery)

**Figure 2:**
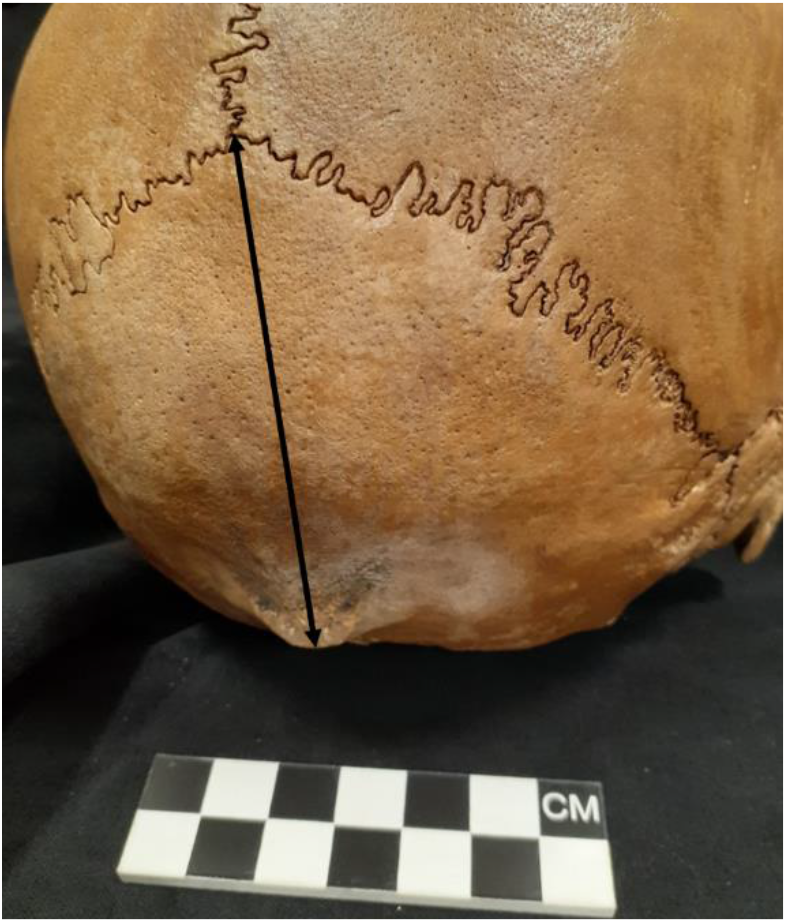
Lambda-Inion distance (Image taken by GT Historic St. Mark’s Military Cemetery)

**Figure 3:**
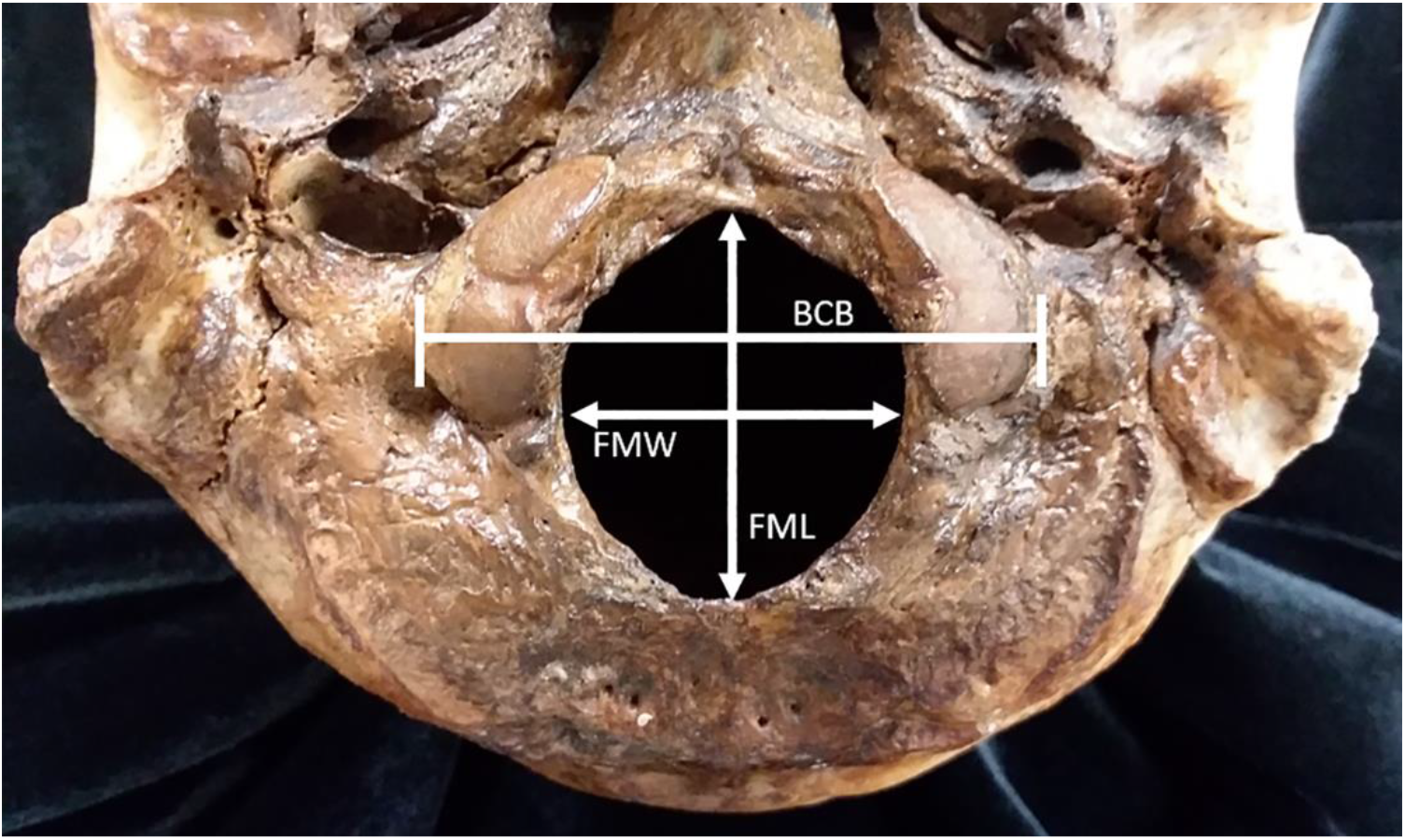
Foramen Magnum Measurements: BCB) Bicondylar breadth in sagittal plane; FMW) foramen magnum breadth in sagittal plane; FML) foramen magnum length in transverse plane. (Image taken by GT Historic St. Mark’s Military Cemetery)

Two trials of data collection were completed (GT) for observations of three nonmetric traits: lateral form of the external occipital protuberance, presence of nuchal crest, and nuchal line count. The lateral form of the external occipital protuberance (Figure 4) is described as flat, round, protuberant, and modified (Olivier, 1969). The presence or absence of a nuchal crest and the nuchal line count(Figure 5) are standard traits (White, 1991).

**Figure 4:**
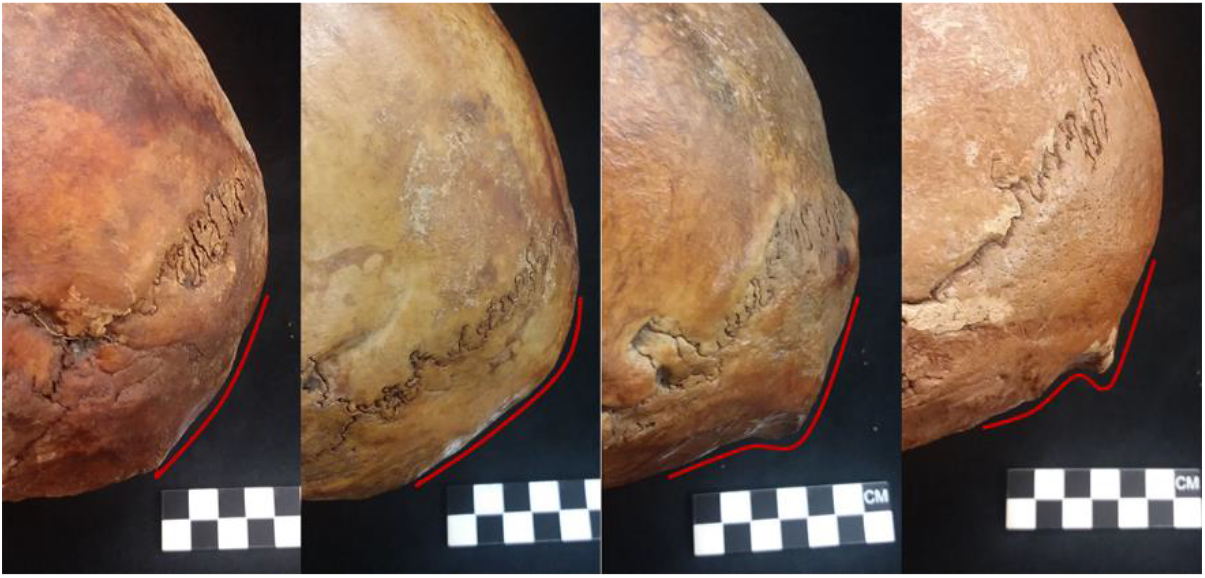
External occipital protuberance general form – Flat, Rounded, Protuberant, and Modified (left to right). (Image taken by GT Historic St. Mark’s Military Cemetery)

**Figure 5:**
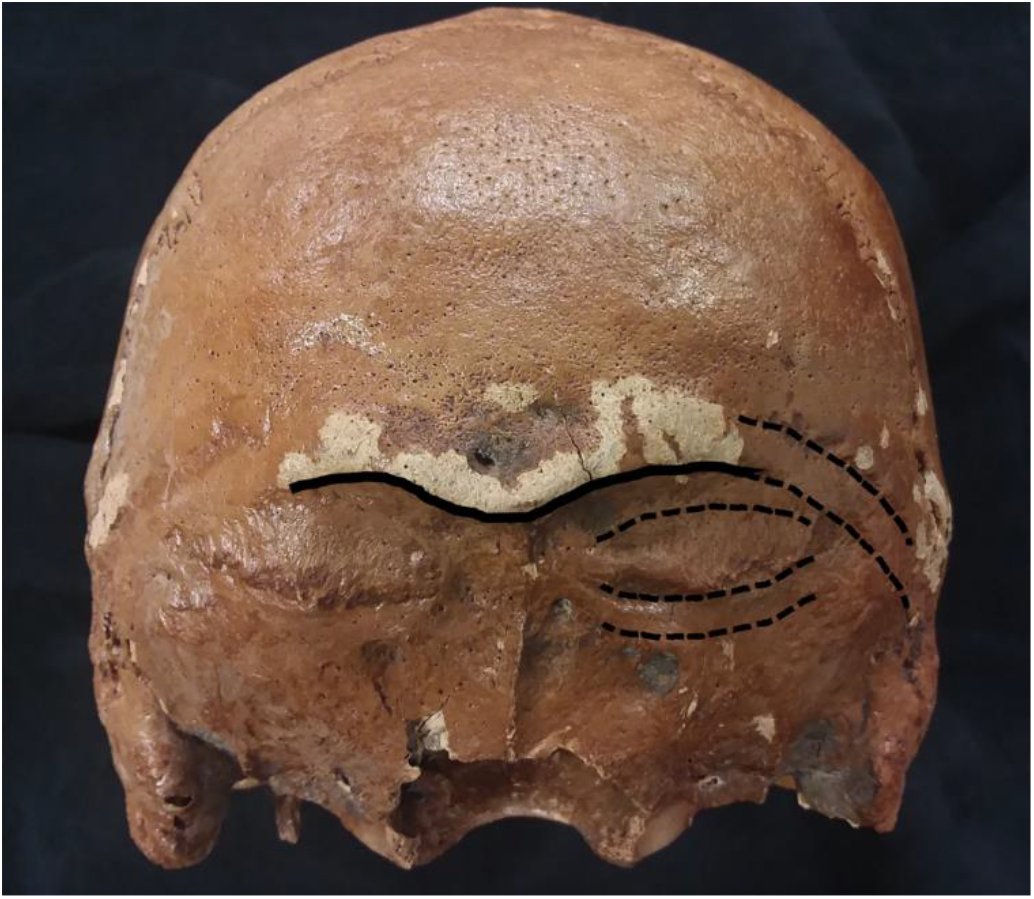
Nuchal crest (solid line) and nuchal lines (dashed line). (Image taken by GT Historic St. Mark’s Military Cemetery)

### Mined Dataset Data Mining

Published data used for this study were not in digital format. The thesis data for the Arikara (Williams, 1987) were in PDF format and OCR was used to identify text. Visual inspection resulted in manual correction of errors introduced by the text recognition process. Thesis data shared by Daniel J. Wescott (Wescott, 1996) were in jpg format. JPGs were cropped to limit data required for this project. Adobe was used to enhance cropped camera images and OCR was used to recognize text. Visual inspection of original and scanned data resulted in manual correction of errors introduced by the conversion process. These data are not analyzed independently but are combined with our data (see next).

### Combined Dataset

For analytical purposes, we combined the Original and Mined Datasets where they overlapped. Variables in this dataset are limited to the foramen magnum and bicondylar breadth measures.

### Statistical Analysis

Data were analyzed (KCH) in R version 4.1.2 “Bird Hippie” (RStudio Team, 2020) using RStudio Desktop 2022.07.0+548 (R Development Core Team, 2018). Data were manipulated using *tidyverse* (Wickham et al., 2019). Agreement for nonmetric traits in the Original Dataset was conducted using the *kappa* function in *irr* (Gamer, Lemon, Fellows, & Singh, 2012). Equality of variance in measurement error was tested for the three trails of data collected from Florida skeletal assemblages using base R functions. Collinearity of variables was tested using the *pairs.panels* function in *psych* (Revelle, 2018). PCA was conducted using base R. Prior to ANOVA, we tested our data met the assumptions of normality using the *qqp* function and variance equality using *leveneTest* in *car* (Fox & Weisberg, 2011) and for outliers using *RosnerTest* in *EnvStats* (Millard, 2013). Descriptive statistics were generated using the *describe.by* function in *psych* (Revelle, 2018). If assumptions were not violated, we used the *lm* function in base R to test for interaction between subsistence and sex. In the absence of interaction, we used the *Anova* function with Type II Sum of Squares in *car* to test model fit for each variable (Fox & Weisberg, 2011). If normality assumptions were violated, *glm* is substituted for *lm* (Fox, 2008). If variance assumption is violated, the white.adjust parameter in *Anova* is used, which tests a heteroscedasticity-corrected coefficient covariance matrix (Fox, 2008; Fox & Weisberg, 2011). Outliers will be examined for fit and removed if extreme values. We used discriminant function analysis to test foramen magnum area and bicondylar breadth accuracy in biological sex estimation using the Combined Dataset using the *lda* function in *MASS* (Venables & Ripley, 2002). All plots were made using *ggplot2* (Wickham, 2016) and *ggpubr* (Kassambara, 2018) and panels were created using *patchwork* (Pedersen, 2020).

### Data Sharing

All data used in this study and scripts for their manipulation and analysis are shared via: https://github.com/kchoover14/OccipitalActivityPatterns

## RESULTS

### Intra-observer Error on Original Dataset

Intra-observer agreement for nonmetric data was assessed using the kappa2 function in the *irr* package (Gamer et al., 2012). There were no significant disagreements between observations for each of the three traits (Table 2). The data for second trial were used. Levene’s statistic was used to assess measurement error for metric traits across three trails using the *car* package (Fox, 2008). There was no significant variance attributable to measurement error (Table 2). We also examined how much measurement error was contributing to trait variable and it was low ≤0.21% (Table 2) and slightly less than in published studies (Wescott & Moore-Jansen, 2001). The average of three measurements was used for hypothesis testing.

**Table 2:** Inter-observer agreement between observations for nonmetric traits and measurement error analysis.

|  |  | **Trials** | **n** | **kappa** | **z-value** | **p-value** |
| --- | --- | --- | --- | --- | --- | --- |
| **Kappa** | Nuchal crest | 2 | 196 | 0.00 | 0.00 | 1.00 |
|  | Nuchal lines | 2 | 196 | 0.00 | 0.00 | 1.00 |
|  | General Form | 2 | 196 | 0.01 | 0.28 | 0.78 |
|  |  | **Trials** | **df1** | **df2** | **F** | **p-value** |
| **Levene's Test** | Bicond Breadth | 3 | 2 | 186 | 0.10 | 0.90 |
|  | Depth EOP | 3 | 2 | 291 | 0.77 | 0.46 |
|  | FM Breadth | 3 | 2 | 189 | 1.68 | 0.19 |
|  | FM Length | 3 | 2 | 201 | 0.63 | 0.54 |
|  | Lambda-Inion | 3 | 2 | 291 | 1.35 | 0.26 |
|  |  | **ME Mean** | **ME SD** | **Trait Mean** | **Trait SD** | **% Error** |
| **Measurement Error** | Bicond Breadth | 0.11 | 0.66 | 51.04 | 3.45 | 0.21% |
|  | Depth EOP | 0.02 | 0.27 | 6.08 | 1.67 | 0.37% |
|  | FM Sagital | 0.00 | 0.27 | 35.98 | 2.57 | 0.01% |
|  | FM Transverse | -0.01 | 0.14 | 30.85 | 2.48 | -0.05% |
|  | Lambda-Inion | 0.33 | 0.14 | 64.43 | 7.18 | 0.51% |

### Dimension Reduction for Original and Combined Datasets

As noted in the introduction, the foramen magnum is under stronger genetic control during early development while the occipital condyles and posterior ectocranial portion of the occipital bone are prone to remodeling in life due to biomechanical processes. To reduce dimensions for statistical modelling, we calculated the foramen magnum ellipse area (pi * length * breadth) to serve as an index (Index_FM) for comparing sex and subsistence groups for differences. Because the bicondylar breadth is the only remaining variable for the skull base, it serves as a contrast to the more genetically controlled foreman magnum area and allows us to explore variation is attributable to each. For the ectocranial portion of the occipital bone subject to biomechanical pressure, we used measures of collinearity and PCA to determine if data reduction was possible. Supplemental Figure 1 (grouped by subsistence) and Supplemental Figure 2 (grouped by sex) each visualize the relationship among variables from the pairs.panel function of psych (Revelle, 2018). The only variable pairs not correlated are lambda-inion distance to nuchal lines and crest. The PCA results indicate that one component explains 50% of the variance but three explain 82% (Supplemental Table 1). Figure 6 visualizes the eigenvectors from the PCA. The vector for the lambda-inion distance diverges from other vectors while depth of the external occipital protuberance and the general form together form one vector and the nuchal lines and crest form another vector. Guided by these results, we reduced data via the creation of two indexes: depth of external occipital protuberance plus general form (Index_Form), nuchal line count plus presence of nuchal crest (Index_Nuchal). The indexes reduce the total dimensions for the Original Dataset from eight to five variables, two for the base of the skull and three for ectocranial occipital bone morphology. The total dimensions of the Combined Dataset are reduced from three to two. Descriptive statistics are found in Supplemental Table 2.

**Figure 6:**
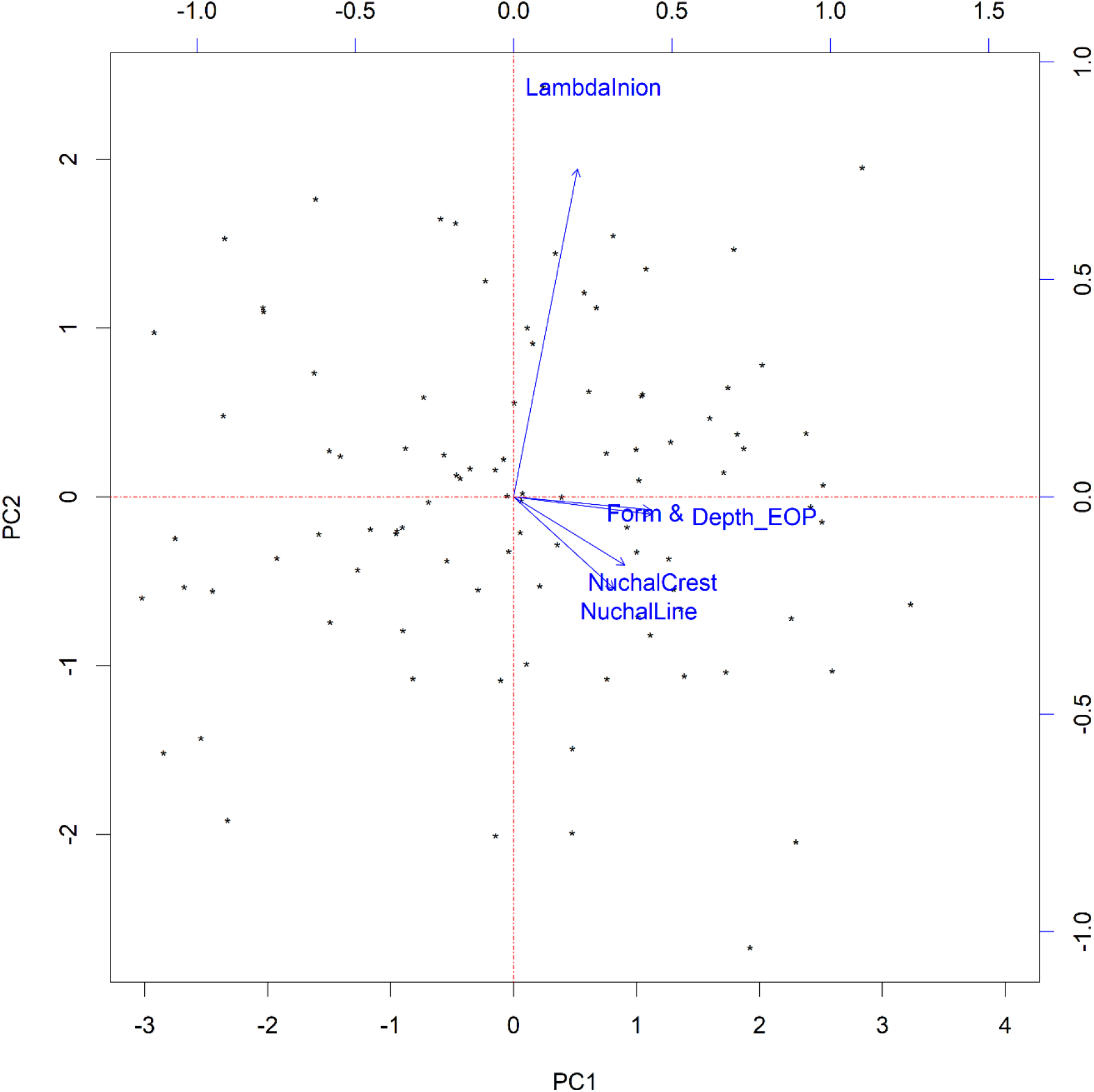
Biplot of PCA eigenvectors for variables subject to biomechanical stress.

**Figure 7:**
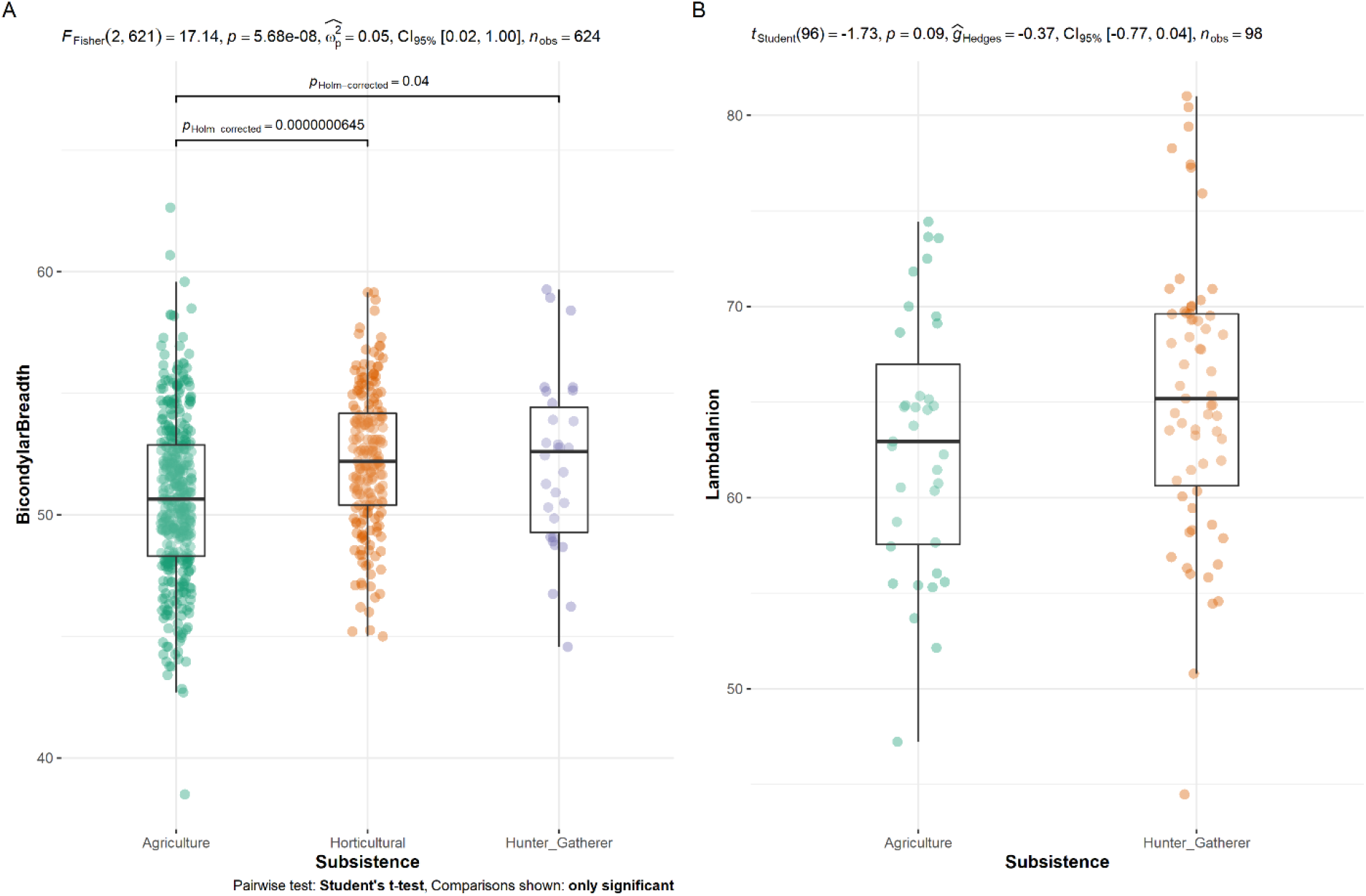
Graphic visualization of significant ANOVA results by subsistence (graph values are for ANOVA, not linear regression values reported in the manuscript). A) Bicondylar Breadth for Combined Dataset; B) Lamba-Inion distance for Original Dataset.

**Figure 8:**
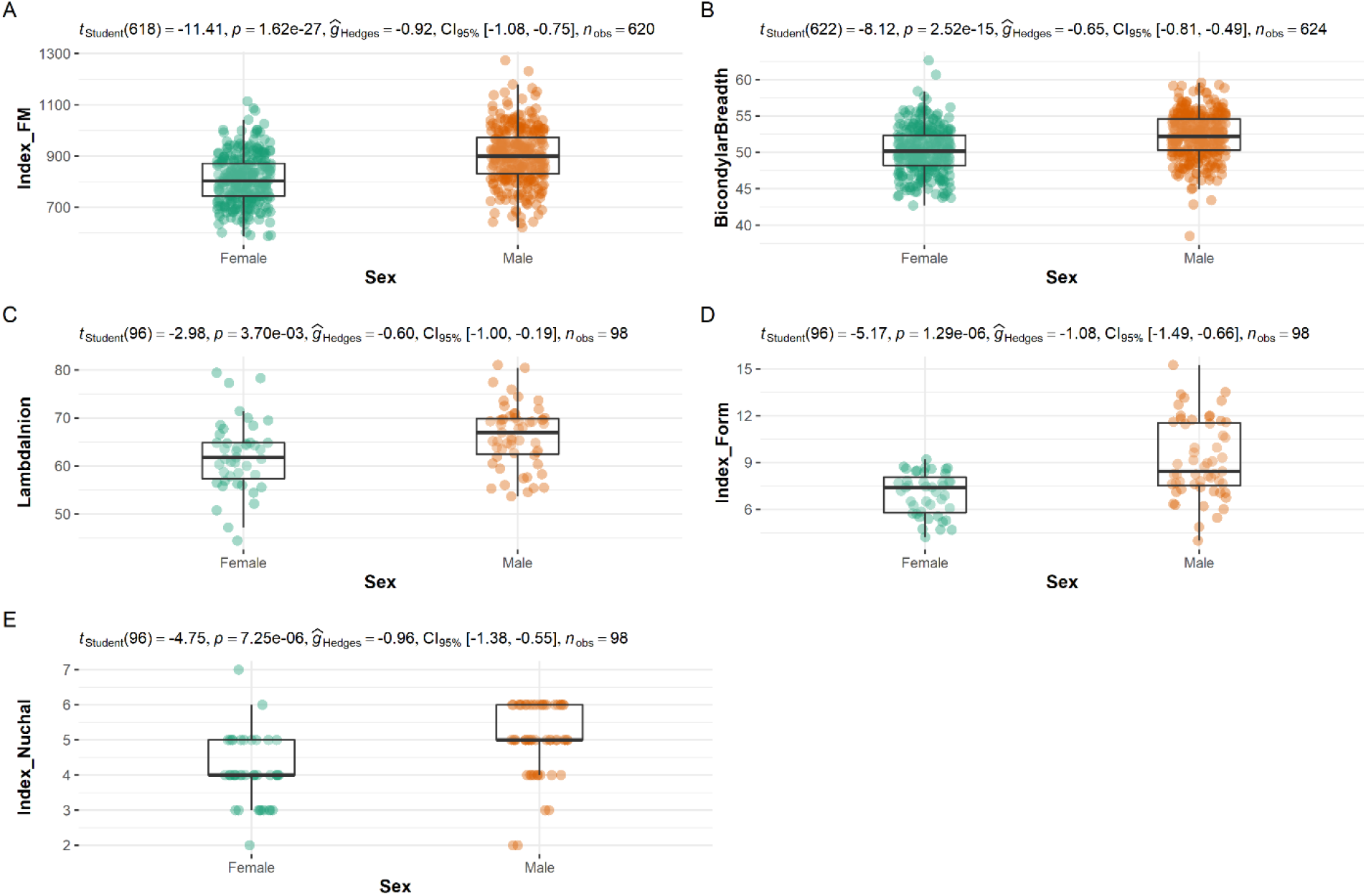
Graphic visualization of significant ANOVA results by sex (graph values are for ANOVA, not linear regression values reported in the manuscript). A) Foramen magnum area for Combined Dataset; B) Bicondylar breadth for Combined Dataset; C) Lamba-Inion distance for Original Dataset; D) Occipital morphology for Original Dataset; E) Nuchal area for Original Dataset.

### Testing ANOVA Assumptions

We tested ANOVA assumptions for the Original Dataset (five variables) and Combined Dataset (two variables) separately. First, we used the *qqp* function in the *car* package to plot empirical quantiles of each against theoretical quantiles of the normal distribution (Fox, 2008). All variables were normally distributed for each dataset with a confidence interval of 95% (based on standard errors of an independent random sample from the comparison distribution (Supplemental Figures 3-7). Second, we used the leveneTest in the *car* package to assess homogeneity of variance across groups (Fox, 2008). In the Original Dataset, the only unequal variance was for Index_Form by sex. In the Combined Dataset the only unequal variance was for Index_FM by sex. See Supplemental Table 3 for results. Third, we used the *RosnerTest* in *EnvStats* (Millard, 2013) to identify outliers. There were no outliers. See Supplemental Table 4 for results.

### Models

We fitted models using subsistence and sex as independent variables and occipital traits as dependent variables. Because unequal variances were found for Index_Form (Original Dataset) and Index_FM (Combined Dataset), we used the white.adjust parameter in Anova for those models (see Methods), which uses least square rather than sum of squares. We first two-way models testing for interaction between dependent variables and found none. In the absence of interaction, two-way (subsistence + sex) ANOVAs with Type II sum of squares method was used (Table 3).

**Table 3:** Two-way ANOVA results.

| Dataset | Variable | Predictor | SS | df | F | p-value |
| --- | --- | --- | --- | --- | --- | --- |
| Original | Index_FM | Subsistence | 41 | 1 | 0.00 | 0.952 |
|  |  | Sex | 103503 | 1 | 9.27 | 0.003 * |
|  |  | Residuals | 680878 | 61 |  |  |
| Combined | Index_FM** | Subsistence |  | 2 | 2.41 | 0.09 |
|  |  | Sex |  | 1 | 129.26 | <0.002 * |
|  |  | Residuals |  | 616 |  |  |
| Original | BicondylarBreadth | Subsistence | 90 | 1 | 9.93 | 0.003 * |
|  |  | Sex | 128 | 1 | 14.12 | <0.002 * |
|  |  | Residuals | 545 | 60 |  |  |
| Combined | BicondylarBreadth | Subsistence | 361 | 2 | 19.6 | <0.002 * |
|  |  | Sex | 654 | 1 | 70.9 | <0.002 * |
|  |  | Residuals | 5719 | 620 |  |  |
| Original | LambdaInion | Subsistence | 267 | 1 | 5.95 | 0.017 * |
|  |  | Sex | 536 | 1 | 11.93 | 0.001 * |
|  |  | Residuals | 4264 | 95 |  |  |
| Original | Index_Form** | Subsistence |  | 1 | 0.18 | 0.670 |
|  |  | Sex |  | 1 | 29.66 | <0.002 * |
|  |  | Residuals | 95 |  |  |  |
| Original | Index_Nuchal | Subsistence | 0.7 | 1 | 0.73 | 0.390 |
|  |  | Sex | 22.2 | 1 | 23.18 | <0.002 * |
|  |  | Residuals | 90.9 | 95 |  |  |

\*Significant value
\*\*Coefficient covariances computed by hccm()

We first examined variables developing early in life and less prone to remodeling later in life. The foramen magnum area is significantly different between the sexes (Table 3), which supports the previously identified sexually dimorphic nature of this variable. The linear model coefficients in Table 4 are in comparison to agricultural values (the most variable of the three subsistence lifestyles) and females. The foramen magnum area increases in size from horticulturalists to agriculturalists to hunter-gatherers, with the most striking difference between horticulturalists and agriculturalists (−18.70 cm^2^). In both datasets, males are larger than females—in the larger combined dataset, the difference in size is ~10 cm^2^. In both datasets, bicondylar breadth, however, subsistence is significantly different among groups after accounting for sex differences (Table 3). Bicondylar breadth increases in size from agriculturalists to hunter-gatherers to horticulturalists.

**Table 4:** Regression coefficients from linear models.

| Dataset | Independent | Pair | Estimate | StErr | R<sup>2</sup> |
| --- | --- | --- | --- | --- | --- |
| Original | Index_FM | Hunter-Gather | 1.61 | 26.74 | 0.134 |
|  |  | Male | 82.86 | 27.21 |  |
| Combined | Index_FM* | Horticultural | -18.70 | -36.06 | 0.180 |
|  |  | Hunter_Gatherer | 0.96 | -36.23 |  |
|  |  | Male | 92.79 | 76.78 |  |
| Original | BicondylarBreadth | Hunter_Gatherer | 2.42 | 0.77 | 0.261 |
|  |  | Male | 2.94 | 0.78 |  |
| Combined | BicondylarBreadth | Horticultural | 1.61 | 1.09 | 0.150 |
|  |  | Hunter-Gatherer | 1.35 | 0.22 |  |
|  |  | Male | 2.05 | 1.57 |  |
| Original | LambdaInion | Hunter-Gatherer | 3.51 | 1.44 | 0.138 |
|  |  | Male | 4.80 | 1.39 |  |
| Original | Index_Form* | Hunter-Gatherer | 0.20 | 0.45 | 0.219 |
|  |  | Male | 2.21 | 0.43 |  |
| Original | Index_Nuchal | Hunter-Gatherer | 0.18 | 0.21 | 0.196 |
|  |  | Male | 0.98 | 0.20 |  |

\*Coefficient covariances computed by hccm()

For the variables potentially prone to remodeling after development (Original Dataset only), only the lambda-inion length was significant different for both sex and subsistence. Hunter-gatherers have a longer lambda-inion distance, 3.51 cm longer than agriculturalists. Males have a longer length (4.8 cm) than females. The two variables describing the muscular development of the occipital were the combined index for nuchal line count and nuchal crest and the combined index for general form (nominal) and depth of the EOP). Both traits were significantly different between the sexes but not among subsistence practices. The index for males is larger than females by 2.21 and the nuchal index increases by 1 in males—either one line or the presence of a nuchal crest, which is coded as 1. Differences indicate greater muscle marking in males.

For the combined dataset with three types of subsistence, Tukey’s HSD Test for multiple comparisons (Table 5) confirms the foramen magnum area coefficient trends—the difference between hunter-gathers and horticulturalists (−0.1832) is negligible compared to the statistically significant differences between agriculturalists and each of the other two subsistence types. Bicondylar breadth shows a different pattern with the only significant different between horticulturalists (the largest foramen magnum mean area) and agriculturalists (the smallest foreman magnum mean area).

**Table 4:**
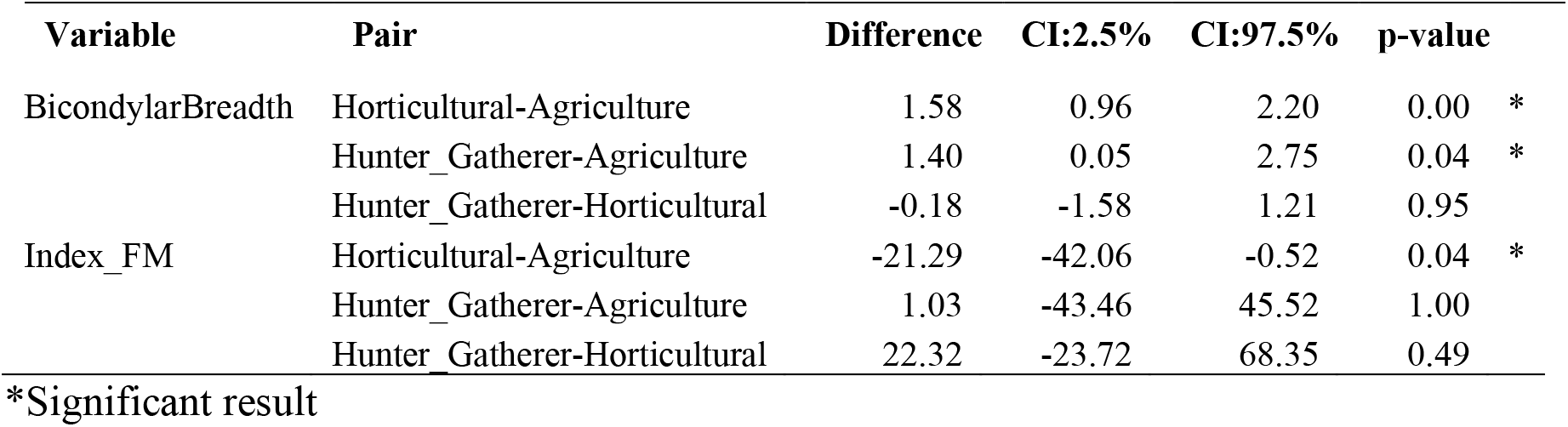
Results of Tukey’s HSD test to determine significant differences between groups.

| Variable | Pair | Difference | CI:2.5% | CI:97.5% | p-value |
| --- | --- | --- | --- | --- | --- |

|  |  |  |  |  |  |  |
| --- | --- | --- | --- | --- | --- | --- |
| BicondylarBreadth | Horticultural-Agriculture | 1.58 | 0.96 | 2.20 | 0.00 | * |
|  | Hunter_Gatherer-Agriculture | 1.40 | 0.05 | 2.75 | 0.04 | * |
|  | Hunter_Gatherer-Horticultural | -0.18 | -1.58 | 1.21 | 0.95 |  |
| Index_FM | Horticultural-Agriculture | -21.29 | -42.06 | -0.52 | 0.04 | * |
|  | Hunter_Gatherer-Agriculture | 1.03 | -43.46 | 45.52 | 1.00 |  |
|  | Hunter_Gatherer-Horticultural | 22.32 | -23.72 | 68.35 | 0.49 |  |

\*Significant result

### Discriminant Function Analysis

We tested the utility of the foramen magnum area in predicting sex. The data for the combined sample were scaled to meet the assumptions of the test and then split into a training set of 70% of the sample and a test set of 30% of the sample. The training set had a female to male ratio of 52:48. The coefficient was 1.108. When applied to the test set, the accuracy in prediction was 71%. When using foramen magnum area and bicondylar breadth, the training set had a 48:52 female to male ratio, with coefficients of 0.89 for foramen magnum area and 0.41 for bicondylar breadth, and an accuracy of 66%.

## DISCUSSION

We examined two areas of the occipital bone, the basicranium and the external posterior of the bone. The basicranium is fully developed by age 7 and contains two areas of interest, the foramen magnum and occipital condyles. Females tend to be smaller than males in basicranium metrics (Catalina-Herrera, 1987; Gangrade et al., 2013; Holland, 1986; Kunar & Nagar, 2014; Macaluso, 2011; Sholapurkar et al., 2017; Zdilla et al., 2017), which may be due to males having larger and heavier heads (Zdilla et al., 2017; S. Zuckerman, 1955). The foramen magnum does not remodel later in life (Scheuer & Black, 2000) sex-based variation in this region reflects early life developmental trajectory. The condyles may remodel later in life under biomechanical pressure from skull rotation, flexion, and extension and sex-based differences in this region (Gangrade et al., 2013; Holland, 1986; Kunar & Nagar, 2014; Macaluso, 2011; Sholapurkar et al., 2017) may be a function of activity rather than inherent sex-based difference (Zdilla et al., 2017; S. Zuckerman, 1955). While the foramen magnum has proven useful in sex estimation of skeletal remains (Gapert & Last, 2008; Gulekon & Turgut, 2003; Hsiao et al., 1996; Sholapurkar et al., 2017; Wescott & Moore-Jansen, 2001), occipital condylar metrics have not been as reliable (Gapert et al., 2009; Wescott & Moore-Jansen, 2001). The posterior portion of the occipital exhibits muscle attachment sites along the nuchal region for head movement and the resulting thickness of the bone and a higher preservation rate has made it a target of study for sex estimation (Gapert & Last, 2008; Gulekon & Turgut, 2003; Hsiao et al., 1996; Sholapurkar et al., 2017; Wescott & Moore-Jansen, 2001; White, 1991). We collected and mined data for five metric traits and three nonmetric traits across human remains practicing hunting-gathering, horticulture, and agriculture. We expected the basicranium to have sex-based differences but no subsistence-based differences. Given the greater influence of biomechanics on the external region of the bone, we expected both sex and subsistence based different in external metric and nonmetric traits. New to this study is the examination of the nuchal area for dimorphism and the use of a foramen magnum index describing the area of the ellipse—the latter reduces variables required for analysis.

We found that the foramen magnum area is only significantly different between the sexes but not subsistence regimes, supporting previous studies and meeting our expectations. The R^2^ values for the models (original and combined datasets) indicated that only 13% of the variance was explained by sex in the Original Dataset and 18% in the Combined Dataset, indicating a low predictive value. We trained a model on a subset of the Combined Dataset (30% of the sample) for discriminant function analysis to determine how accurate sex could be predicted based on foramen magnum area. When tested with the remaining 70% of the Combined Dataset, the accuracy was 71%, which is good but not sufficient for forensic settings.

Bicondylar breadth was significantly different between sexes and among subsistence practices. As noted above, condylar metrics have exhibited sex-based variation in previous studies (Gangrade et al., 2013; Holland, 1986; Kunar & Nagar, 2014; Macaluso, 2011; Sholapurkar et al., 2017). While condyle size might be sexual dimorphism as a function of cranial size (Zdilla et al., 2017; S. Zuckerman, 1955), the variation attributable to subsistence is likely from biomechanical processes (Uysal et al., 2005). Because occipital condylar metrics do not consistently estimate sex accurately (Gapert et al., 2009; Wescott & Moore-Jansen, 2001), perhaps gendered division of labor influences variation along with cranial size and that noise in the signal prevents the trait from being consistent. The slightly larger mean values in hunter-gatherers and horticulturalists (both having a gender-based division of labor) may reflect more physically intense lifestyles compared to the diverse set of occupations associated with agricultural populations. For the Combined Dataset the R^2^ for this model about the same as models for the foramen magnum area alone, 15%. For the Original Dataset, the R^2^ was much higher at 26%. Using both bicondylar breadth and foramen magnum for the discriminant function analysis, resulted in a decline in accuracy from 71% to 66% further supporting the confounding role of biomechanics in bicondylar breadth metrics.

The external area of the occipital region is not frequently examined and we were not able to find published data for this area, limiting our analysis to the Original Dataset that we collected. All traits in this area exhibited sex-based differences. Males had deeper external occipital protuberances, longer lambda-inion distances, and a rougher appearance of the region (including higher nuchal line counts and nuchal crest presence). Our expectation that this area might exhibit variation due to biomechanical activity from subsistence was not met other than for the distance from the lambda to the inion. The R^2^ for these models was. The lambda-inion distance has previously been found to be longer in males (Olivier, 1975) and our data support that finding. We also found, however, that the lambda-inion measure differs by subsistence practice with greater distances in hunter-gatherers. The R^2^ for the models varied with 14% of variance explained by lambda-inion distance, 22% by Index_Form, and 20% by Index_Nuchal. The Original Dataset is too small to split into training and test sets to produce robust discriminant analysis tests, but these traits might have utility in sex estimation and warrant further investigation in larger datasets. The only caveat is that the lambda-inion may be a function of external occipital protuberance morphology and could be duplicative.

Our study took a few novel approaches: the examination of foramen magnum area (rather than transverse and sagittal measurements), the inclusion of metric and nonmetric traits on the ectocranial area of the occipital bone, and the use of training and testing datasets for predicting sex based on occipital traits. We aimed to study the influence of activity on the morphology of the bone by using a diverse set of materials so that we could understand what factors other than sex might be shaping this bone, given the lack of consensus in prior work indicating its utility in forensic settings. Our analysis identified only two variables that had significant subsistence based differences, bicondylar breadth and lambda-inion distance, which suggests possible contributions to variation from biomechanical activity. The occipital bone exhibits sexual dimorphism but the accuracy of sex estimation is not sufficiently high and the use of such metrics and nonmetrics may result in inconsistent classification. It may be supportive alongside other estimates of sex but not on its own. The unexplained variation in the bone does not appear to be strongly influenced by biomechanical activity due to subsistence but other activity patterns may influence variation and the question remains open.

## Acknowledgements

The authors would like to thank the Bureau of Archaeological Research collections manager (Sam Wilford) for allowing us access to and providing us assistance with the Hutchinson Island site materials and Frank L. Williams at Georgia State University for his work in developing the early concepts and baseline for this publication.

