## Supplemental Information for "Sexual dimorphism and biomechanical loading in occipital bone morphological variation"

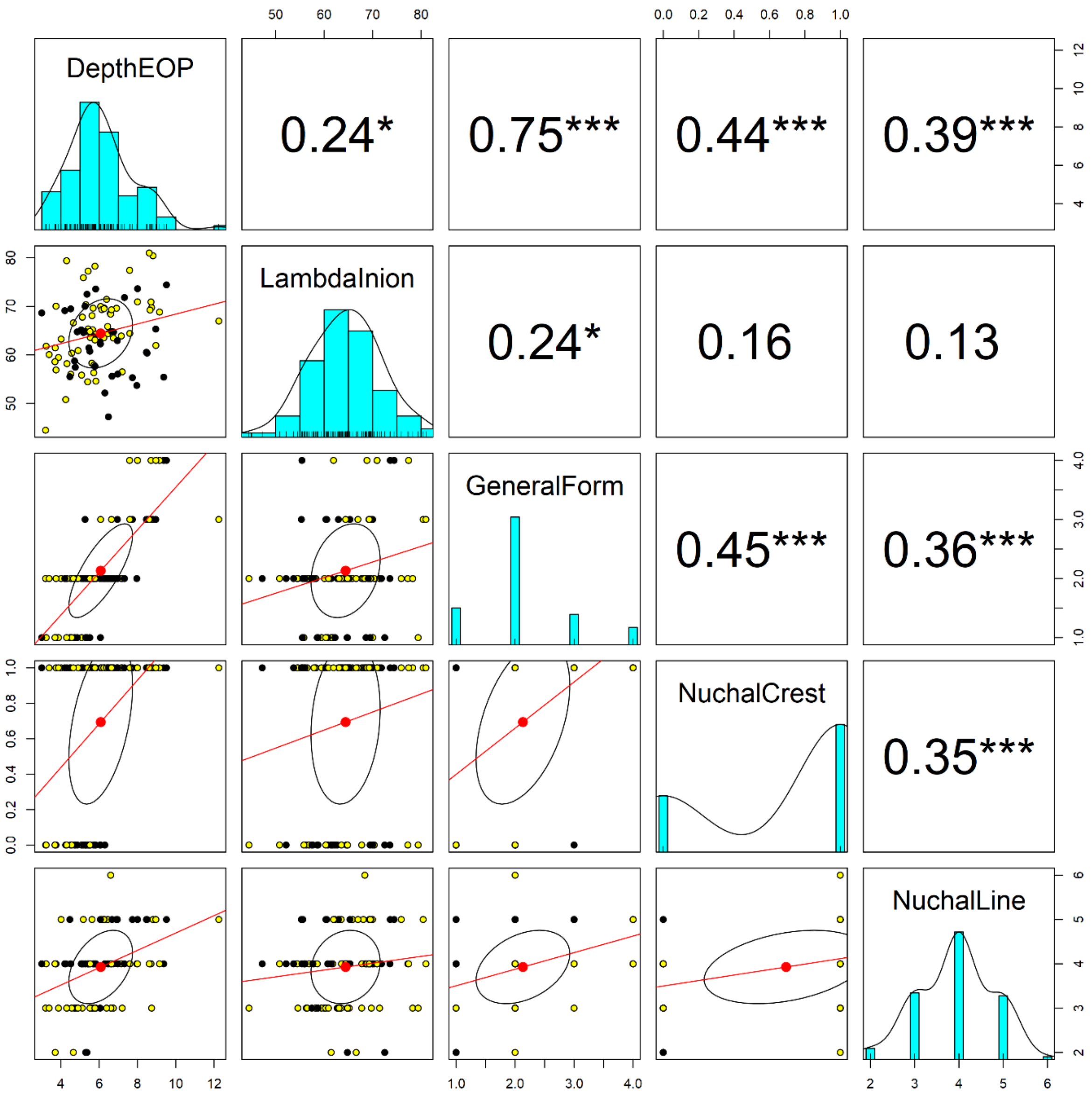

**Supplemental Figure 1: Scatter plot of matrices by subsistence for Original Dataset (black, agriculture; yellow, hunting-gathering. Bivariate scatter plots appear below the diagonal, histograms on the diagonal, and Pearson correlation above the diagonal.**

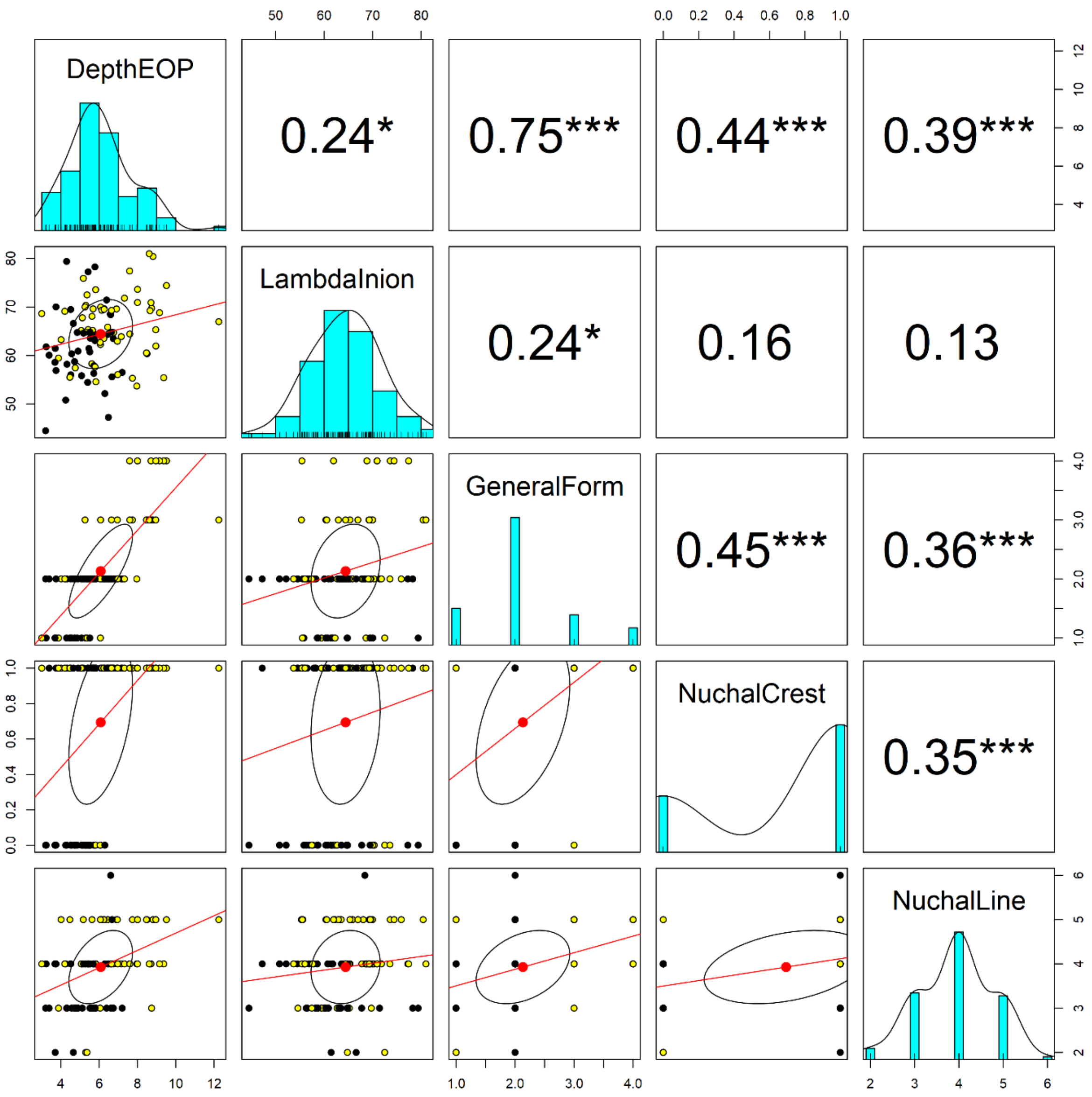

**Supplemental Figure 2: Scatter plot of matrices by sex for Original Dataset (black, Female; yellow, Male). Bivariate scatter plots appear below the diagonal, histograms on the diagonal, and Pearson correlation above the diagonal.**

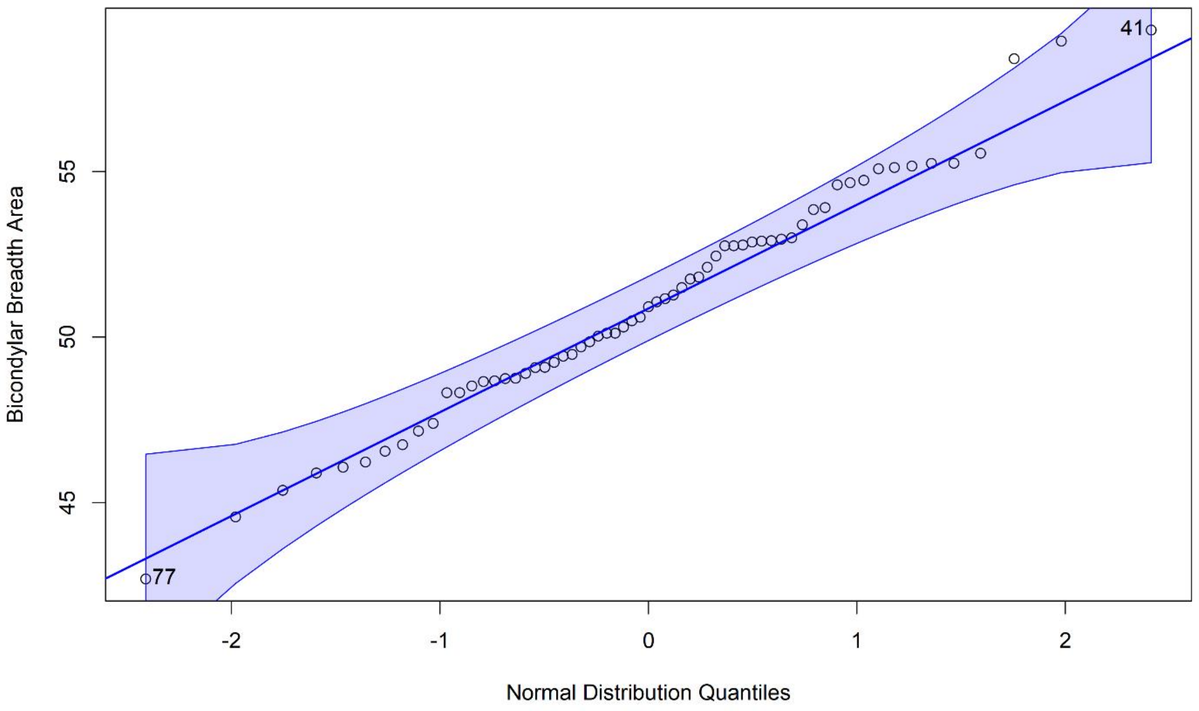

**Supplemental Figure 3: Testing the normal distribution for bicondylar breadth, Original Dataset**

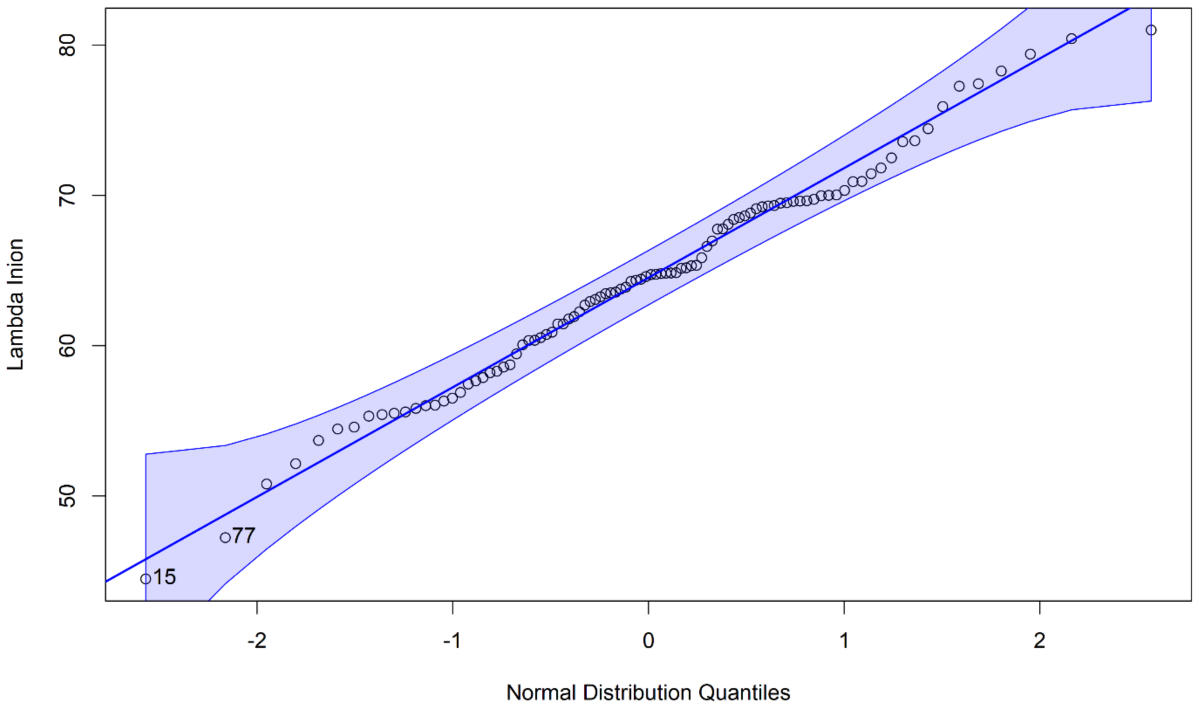

**Supplemental Figure 4: Testing the normal distribution for lambda-inion, Original Dataset**

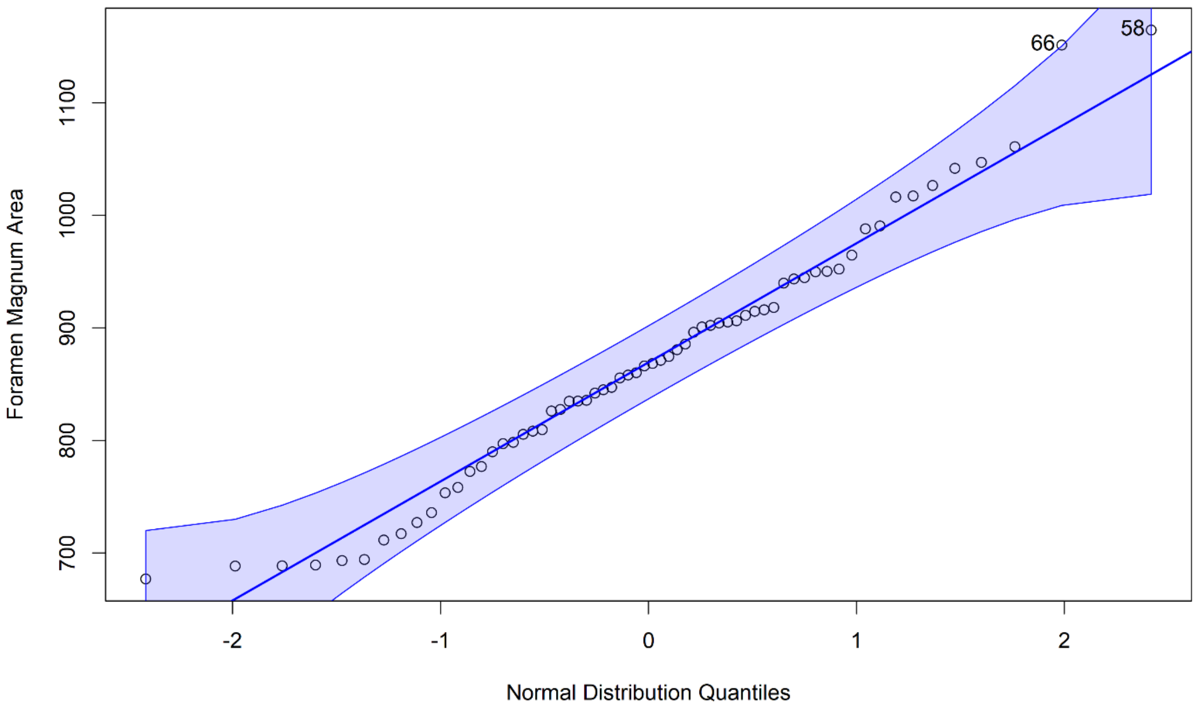

**Supplemental Figure 5: Testing the normal distribution for Index_FM, Original Dataset**

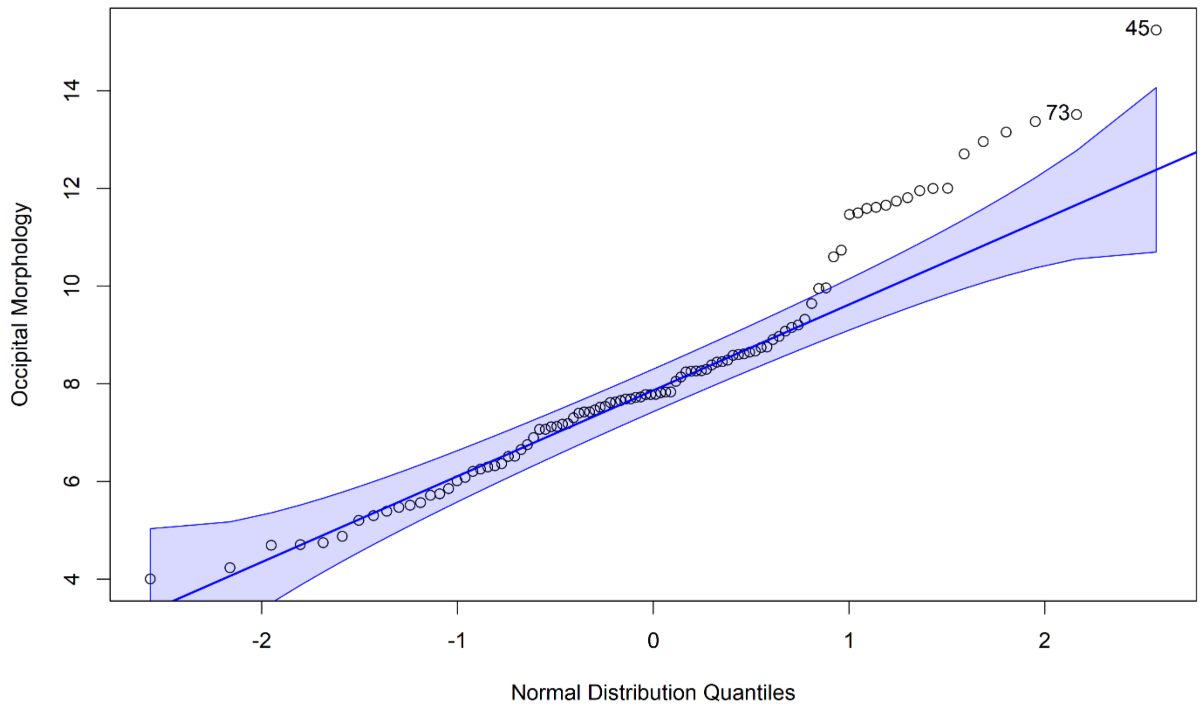

**Supplemental Figure 6: Testing the normal distribution for Index_Form, Original Dataset**

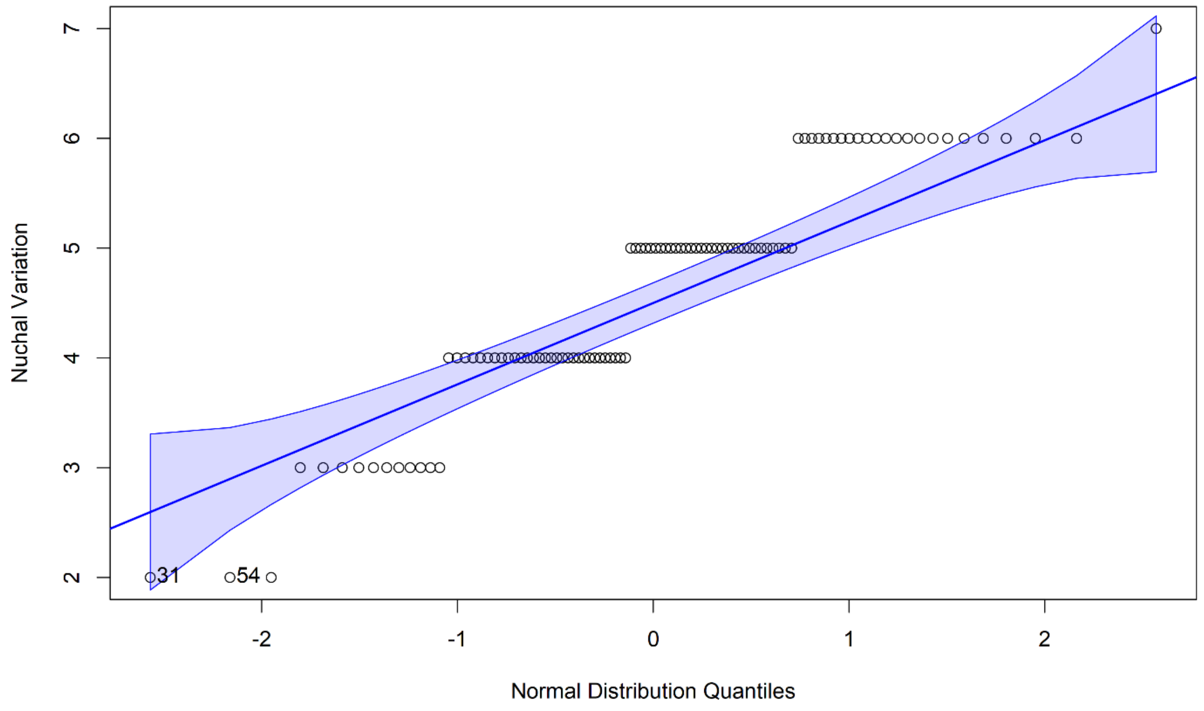

**Supplemental Figure 7: Testing the normal distribution for Index_Nuchal, Original Dataset**

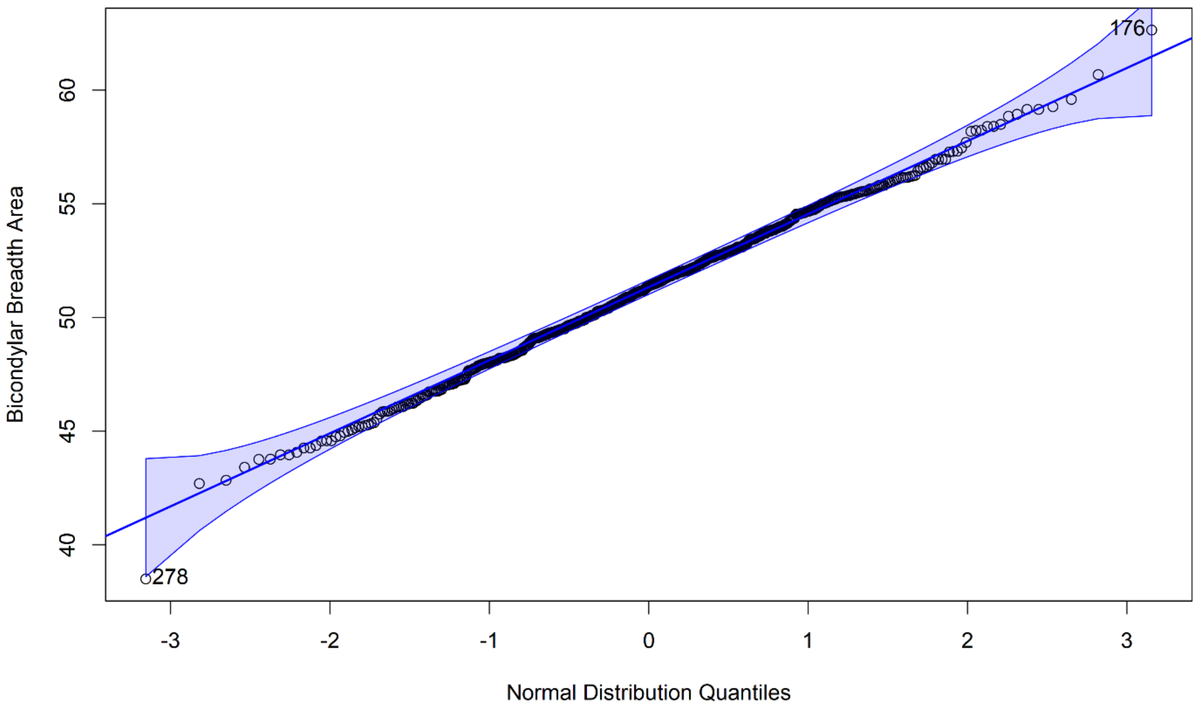

**Supplemental Figure 8: Testing the normal distribution for bincondylar breadth, Combined Dataset**

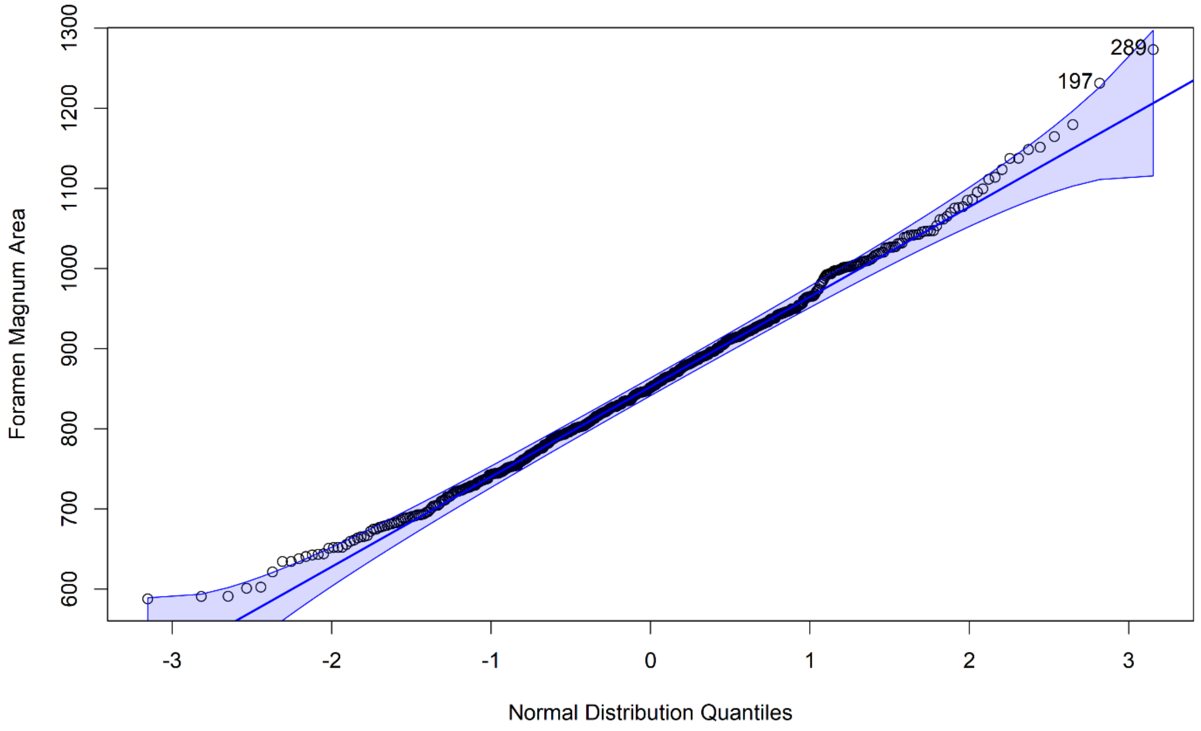

**Supplemental Figure 9: Testing the normal distribution for Index_FM, Combined Dataset**

**Supplemental Table 1: PCA Results**

| | PC1 | PC2 | PC3 | PC4 | PC5 |
|---|---|---|---|---|---|
| DepthEOP | 0.5444 | -0.0506 | 0.3486 | -0.2691 | -0.7121 |
| LambdaInion | 0.2522 | 0.9426 | -0.2088 | 0.0664 | -0.0015 |
| GeneralForm | 0.5394 | -0.0371 | 0.4064 | -0.2284 | 0.7002 |
| NuchalCrest | 0.4390 | -0.1963 | -0.0779 | 0.8731 | -0.0185 |
| NuchalLine | 0.3954 | -0.2630 | -0.8147 | -0.3296 | 0.0468 |
| Variance Explained | 50.00% | 18.16% | 14.39% | 12.56% | 4.88% |

### Supplemental Table 2: Descriptive Statistics

| Dataset | Subsistence | Variable | n | Mean | StDev | StErr | Median | Range |
|---|---|---|---|---|---|---|---|---|
| Original | Hunter-Gatherer | BicondylarBreadth | 25 | 49.46 | 3.33 | 0.67 | 49.09 | 12.55 |
| | | LambdaInion | 43 | 62.09 | 7.37 | 1.12 | 61.78 | 34.93 |
| | | Index_FM | 26 | 820.17 | 83.38 | 16.35 | 820.11 | 299.49 |
| | | Index_Form | 43 | 6.99 | 1.35 | 0.21 | 7.40 | 4.97 |
| | | Index_Nuchal | 43 | 4.09 | 0.92 | 0.14 | 4.00 | 5.00 |
| | Agriculture | BicondylarBreadth | 33 | 50.07 | 3.05 | 0.53 | 50.03 | 12.85 |
| | | LambdaInion | 35 | 62.77 | 6.72 | 1.14 | 62.94 | 27.21 |
| | | Index_FM | 33 | 874.49 | 127.87 | 22.26 | 874.60 | 487.78 |
| | | Index_Form | 35 | 8.36 | 2.33 | 0.39 | 8.05 | 9.51 |
| | | Index_Nuchal | 35 | 4.63 | 1.17 | 0.20 | 5.00 | 4.00 |
| | Female | BicondylarBreadth | 25 | 49.46 | 3.33 | 0.67 | 49.09 | 12.55 |
| | | LambdaInion | 43 | 62.09 | 7.37 | 1.12 | 61.78 | 34.93 |
| | | Index_FM | 26 | 820.17 | 83.38 | 16.35 | 820.11 | 299.49 |
| | | Index_Form | 43 | 6.99 | 1.35 | 0.21 | 7.40 | 4.97 |
| | | Index_Nuchal | 43 | 4.09 | 0.92 | 0.14 | 4.00 | 5.00 |
| | Male | BicondylarBreadth | 38 | 52.07 | 3.16 | 0.51 | 51.93 | 12.72 |
| | | LambdaInion | 55 | 66.25 | 6.46 | 0.87 | 66.97 | 27.31 |
| | | Index_FM | 38 | 902.78 | 117.07 | 18.99 | 901.53 | 487.78 |
| | | Index_Form | 55 | 9.17 | 2.48 | 0.33 | 8.44 | 11.24 |
| | | Index_Nuchal | 55 | 5.04 | 1.02 | 0.14 | 5.00 | 4.00 |
| Combined | Hunter-Gatherer | BicondylarBreadth | 30 | 52.10 | 3.60 | 0.66 | 52.60 | 14.70 |
| | | Index_FM | 31 | 863.60 | 93.30 | 16.76 | 860.20 | 352.50 |
| | Horticultural | BicondylarBreadth | 199 | 52.28 | 2.87 | 0.20 | 52.20 | 14.15 |
| | | Index_FM | 199 | 841.31 | 110.62 | 7.84 | 830.00 | 588.77 |
| | Agriculture | BicondylarBreadth | 395 | 50.70 | 3.33 | 0.17 | 50.65 | 24.14 |
| | | Index_FM | 390 | 862.60 | 113.30 | 5.74 | 860.27 | 685.41 |
| | Female | BicondylarBreadth | 307 | 50.24 | 3.14 | 0.18 | 50.17 | 19.94 |
| | | Index_FM | 306 | 808.59 | 94.17 | 5.38 | 802.64 | 526.06 |
| | Male | BicondylarBreadth | 317 | 52.27 | 3.11 | 0.17 | 52.20 | 21.09 |
| | | Index_FM | 314 | 901.84 | 108.54 | 6.12 | 899.30 | 651.81 |

**Supplemental Table 3: ANOVA Assumptions: Levene's Test for equality of variance**

| Data | Variable | Level | df1 | df2 | F | p-value | |
|---|---|---|---|---|---|---|---|
| Original | BicondylarBreadth | Subsistence | 1 | 61 | 0.69 | 0.410 | |
| | | Sex | 1 | 61 | 0.03 | 0.860 | |
| | LambdaInion | Subsistence | 1 | 96 | 0.08 | 0.780 | |
| | | Sex | 1 | 96 | 0.17 | 0.680 | |
| | Index_FM | Subsistence | 1 | 62 | 2.08 | 0.015 | |
| | | Sex | 1 | 62 | 1.44 | 0.230 | |
| | Index_Form | Subsistence | 1 | 96 | 0.15 | 0.700 | |
| | | Sex | 1 | 96 | 10.80 | 0.001 | * |
| | Index_Nuchal | Subsistence | 1 | 96 | 0.03 | 0.860 | |
| | | Sex | 1 | 96 | 0.74 | 0.392 | |
| Combined | BicondylarBreadth | Subsistence | 2 | 621 | 2.56 | 0.078 | |
| | | Sex | 1 | 622 | 0.08 | 0.780 | |
| | Index_FM | Subsistence | 2 | 617 | 0.79 | 0.460 | |
| | | Sex | 1 | 618 | 3.99 | 0.046 | * |

***Significant result**

### Supplemental Table 4: ANOVA Assumptions: Rosner's Test for multiple outliers

| Data | Variable | n | i | Mean.i | SD.i | Value | Obs.Num | R.i+1 | lambda.i+1 | Outlier |
|---|---|---|---|---|---|---|---|---|---|---|
| Original | BicondylarBreadth | 63 | 0 | 51.04 | 3.45 | 42.70 | 77.00 | 2.42 | 3.22 | FALSE |
| | | | 1 | 51.17 | 3.31 | 59.27 | 41.00 | 2.45 | 3.21 | FALSE |
| | | | 2 | 51.04 | 3.16 | 58.94 | 39.00 | 2.50 | 3.21 | FALSE |
| | | | 3 | 50.91 | 3.02 | 58.40 | 10.00 | 2.49 | 3.20 | FALSE |
| | | | 4 | 50.78 | 2.88 | 44.57 | 4.00 | 2.16 | 3.19 | FALSE |
| | LambdaInion | 98 | 0 | 64.43 | 7.14 | 44.48 | 15.00 | 2.79 | 3.38 | FALSE |
| | | | 1 | 64.63 | 6.88 | 47.23 | 77.00 | 2.53 | 3.37 | FALSE |
| | | | 2 | 64.81 | 6.68 | 81.00 | 96.00 | 2.42 | 3.37 | FALSE |
| | | | 3 | 64.64 | 6.50 | 80.43 | 17.00 | 2.43 | 3.37 | FALSE |
| | | | 4 | 64.47 | 6.33 | 79.40 | 92.00 | 2.36 | 3.36 | FALSE |
| | Index_FM | 64 | 0 | 869.20 | 111.72 | 1165.00 | 58.00 | 2.65 | 3.22 | FALSE |
| | | | 1 | 864.50 | 106.07 | 1151.00 | 66.00 | 2.70 | 3.22 | FALSE |
| | | | 2 | 859.90 | 100.33 | 1061.00 | 56.00 | 2.01 | 3.21 | FALSE |
| | | | 3 | 856.60 | 97.71 | 1047.00 | 55.00 | 1.95 | 3.21 | FALSE |
| | | | 4 | 853.40 | 95.31 | 1042.00 | 39.00 | 1.98 | 3.20 | FALSE |
| | Index_Form | 98 | 0 | 8.21 | 2.32 | 15.24 | 45.00 | 3.03 | 3.38 | FALSE |
| | | | 1 | 8.14 | 2.22 | 13.51 | 73.00 | 2.42 | 3.37 | FALSE |
| | | | 2 | 8.09 | 2.16 | 13.37 | 75.00 | 2.44 | 3.37 | FALSE |
| | | | 3 | 8.03 | 2.10 | 13.15 | 88.00 | 2.44 | 3.37 | FALSE |
| | | | 4 | 7.98 | 2.05 | 12.96 | 97.00 | 2.44 | 3.36 | FALSE |
| | Index_Nuchal | 98 | 0 | 4.62 | 1.08 | 2.00 | 31.00 | 2.43 | 3.38 | FALSE |
| | | | 1 | 4.65 | 1.05 | 2.00 | 54.00 | 2.52 | 3.37 | FALSE |
| | | | 2 | 4.68 | 1.02 | 2.00 | 55.00 | 2.62 | 3.37 | FALSE |
| | | | 3 | 4.71 | 0.99 | 7.00 | 37.00 | 2.32 | 3.37 | FALSE |
| | | | 4 | 4.68 | 0.96 | 3.00 | 2.00 | 1.74 | 3.36 | FALSE |
| Combined | BicondylarBreadth | 624 | 0 | 51.27 | 3.285 | 38.5 | 278 | 3.89 | 3.921 | FALSE |
| | | | 1 | 51.29 | 3.248 | 62.64 | 176 | 3.49 | 3.921 | FALSE |
| | | | 2 | 51.27 | 3.218 | 60.68 | 137 | 2.92 | 3.92 | FALSE |
| | | | 3 | 51.26 | 3.199 | 42.7 | 77 | 2.68 | 3.92 | FALSE |
| | | | 4 | 51.27 | 3.183 | 42.84 | 436 | 2.65 | 3.919 | FALSE |
| | Index_FM | 620 | 0 | 855.8 | 111.8 | 1273 | 289 | 3.73 | 3.919 | FALSE |
| | | | 1 | 855.1 | 110.6 | 1231 | 197 | 3.4 | 3.919 | FALSE |
| | | | 2 | 854.5 | 109.7 | 1180 | 520 | 2.96 | 3.919 | FALSE |
| | | | 3 | 854 | 109 | 1165 | 58 | 2.85 | 3.918 | FALSE |
| | | | 4 | 853.5 | 108.4 | 1151 | 66 | 2.75 | 3.918 | FALSE |
